## Supplementary Material for "Generalized Michaelis–Menten rate law with time-varying molecular concentrations"

The technical details of numerical simulation and statistical analysis methods are presented in Text S12. Equation and figure numbers without "S" refer to those in the main text.

**Text S1. Rate law overview and derivation**

Consider two different molecules A and B that bind to each other and form complex AB. The concentration of the complex AB at time $t$ is denoted by $C(t)$ and its dynamics is governed by Eq. (1). In Eq. (1), $A(t)$ and $B(t)$ denote the total concentrations of A and B, respectively, and their temporal profiles are allowed to be very generic, e.g., even with their own feed-back effects as in the example applications in this study. In Eq. (1), $k_{\mathrm{a}}$ and $k_{\delta}$ are rate param-eters. For the sake of generality, $k_{\delta}$ is not limited to AB's dissociation event but encompasses all rate events to lower the level of AB. Using the notations $\tau \equiv k_{\delta}t$, $K \equiv k_{\delta}/k_{\mathrm{a}}$, $\bar{A}(\tau) \equiv A(t)/K$, $\bar{B}(\tau) \equiv B(t)/K$, and $\bar{C}(\tau) \equiv C(t)/K$, one can rewrite Eq. (1) as

$$\frac{\mathrm{d}\bar{C}(\tau)}{\mathrm{d}\tau} = [\bar{A}(\tau) - \bar{C}(\tau)][\bar{B}(\tau) - \bar{C}(\tau)] - \bar{C}(\tau). \quad \text{(S1)}$$

By definition, $C(t) \leq A(t)$ and $C(t) \leq B(t)$, and therefore

$$C(t) \leq \min[A(t), B(t)] \text{ {i.e., } } \overline{C}(\tau) \leq \min[\overline{A}(\tau), \overline{B}(\tau)]\text{}. \quad \text{(S2)}$$

On the other hand, Eq. (S1) is equivalent to

$$\frac{\mathrm{d}\bar{C}(\tau)}{\mathrm{d}\tau} = \left[\bar{C}(\tau) - \bar{C}_{\mathrm{tQ}}(\tau)\right]\left\{\bar{C}(\tau) - \left[\bar{C}_{\mathrm{tQ}}(\tau) + \Delta_{\mathrm{tQ}}(\tau)\right]\right\}, \quad \text{(S3)}$$

where $\bar{C}_{\mathrm{tQ}}(\tau)$ and $\Delta_{\mathrm{tQ}}(\tau)$ are given by

$$\bar{C}_{\mathrm{tQ}}(\tau) \equiv \frac{1}{2}\left[1 + \overline{A}(\tau) + \overline{B}(\tau) - \Delta_{\mathrm{tQ}}(\tau)\right], \quad \text{(S4)}$$

$$\Delta_{\mathrm{tQ}}(\tau) \equiv \sqrt{[1 + \overline{A}(\tau) + \overline{B}(\tau)]^2 - 4\overline{A}(\tau)\overline{B}(\tau)}. \quad \text{(S5)}$$

Of note, $\bar{C}_{\mathrm{tQ}}(\tau) = C_{\mathrm{tQ}}(t)/K$ and $\Delta_{\mathrm{tQ}}(\tau) = \Delta_{\mathrm{tQ}}(t)$ from the definitions of $C_{\mathrm{tQ}}(t)$ and $\Delta_{\mathrm{tQ}}(t)$ in Eqs. (2) and (3), respectively. In the tQSSA, the assumption is that $C(t)$ approaches an equilibrium (quasi-steady state) fast enough each time, given the values of $A(t)$ and $B(t)$ [S1–S3]. To understand this idea, notice that $\bar{C}'(\tau) \to 0$ in Eq. (S3) when $\bar{C}(\tau) \to \bar{C}_{\mathrm{tQ}}(\tau)$ given the values of $\overline{A}(\tau)$ and $\overline{B}(\tau)$ [we use symbol ′ for a derivative, such as $\bar{C}'(\tau)$ here]. One can prove that $\bar{C}_{\mathrm{tQ}}(\tau) \leq \min[\overline{A}(\tau), \overline{B}(\tau)]$ and thus Eq. (S2) is naturally satisfied when $\bar{C}(\tau) = \bar{C}_{\mathrm{tQ}}(\tau)$. The other nominal solution of $\bar{C}'(\tau) = 0$ in Eq. (S3) does not satisfy Eq. (S2) and is thus physically senseless.

According to the tQSSA, one takes an estimate $\bar{C}(\tau) \approx \bar{C}_{\mathrm{tQ}}(\tau)$, or equivalently, $C(t) \approx C_{\mathrm{tQ}}(t)$. The tQSSA is generally more accurate than the conventional MM rate law [S1–S4]. Under the assumption of Eq. (4), which is essentially identical to the assumption in the sQSSA [S5], the Padé approximant for $\bar{C}_{\mathrm{tQ}}(\tau)$ takes the following form:

$$\bar{C}_{\mathrm{tQ}}(\tau) \approx \frac{\overline{A}(\tau)\overline{B}(\tau)}{1+\overline{A}(\tau)+\overline{B}(\tau)}. \tag{S6}$$

Eq. (S6) is equivalent to Eq. (5). In the example of a typical metabolic reaction with $B(t) \ll A(t)$ for substrate A and enzyme B, Eq. (4) is automatically satisfied and Eq. (S6) further reduces to the MM rate law $\bar{C}_{\mathrm{tQ}}(\tau) \approx \overline{A}(\tau)\overline{B}(\tau)/\left[1+\overline{A}(\tau)\right]$, the outcome of the sQSSA [S1,S5–S9].

Both the above tQSSA and sQSSA stand on the assumption that $C(t)$ approaches the quasi-steady state fast enough each time before the marked temporal change of $A(t)$ or $B(t)$. We here relieve this quasi-steady state assumption and develop the ETS as the better approximation of $C(t)$ in the case of time-varying $A(t)$ and $B(t)$. Suppose that $C(t)$ may not necessarily approach the quasi-steady state $C_{\mathrm{tQ}}(t)$ but stays within some distance from it, satisfying the following relation:

$$\left|\bar{C}(\tau) - \bar{C}_{\mathrm{tQ}}(\tau)\right| \ll \Delta_{\mathrm{tQ}}(\tau). \tag{S7}$$

This relation is readily satisfied in physiologically-relevant conditions (Text S5). This relation allows us to discard $\left[\bar{C}(\tau) - \bar{C}_{\mathrm{tQ}}(\tau)\right]^2$ compared to $\Delta_{\mathrm{tQ}}(\tau)\left[\bar{C}(\tau) - \bar{C}_{\mathrm{tQ}}(\tau)\right]$ and thereby reduce Eq. (S3) to

$$\frac{\mathrm{d}\bar{C}(\tau)}{\mathrm{d}\tau} \approx \Delta_{\mathrm{tQ}}(\tau)\left[\bar{C}_{\mathrm{tQ}}(\tau) - \bar{C}(\tau)\right]. \tag{S8}$$

Multiplying both the left- and right-hand sides above by $e^{\int_{\tau_0}^{\tau}\Delta_{\mathrm{tQ}}(\tau')\mathrm{d}\tau'}$ (where $\tau_0$ denotes an arbitrarily assigned, initial point of $\tau$) and applying the product rule lead to

$$\frac{\mathrm{d}}{\mathrm{d}\tau}\left[\bar{C}(\tau)e^{\int_{\tau_0}^{\tau}\Delta_{\mathrm{tQ}}(\tau')\mathrm{d}\tau'}\right] \approx \Delta_{\mathrm{tQ}}(\tau)\bar{C}_{\mathrm{tQ}}(\tau)e^{\int_{\tau_0}^{\tau}\Delta_{\mathrm{tQ}}(\tau')\mathrm{d}\tau'}. \tag{S9}$$

The integral of Eq. (S9) from $\tau = \tau_0$ ends up the following solution of Eq. (S8):

$$\bar{C}(\tau) \approx \int_{\tau_0}^{\tau}\Delta_{\mathrm{tQ}}(\tau')\bar{C}_{\mathrm{tQ}}(\tau')e^{-\int_{\tau'}^{\tau}\Delta_{\mathrm{tQ}}(\tau'')\mathrm{d}\tau''}\mathrm{d}\tau' + \bar{C}(\tau_0)e^{-\int_{\tau_0}^{\tau}\Delta_{\mathrm{tQ}}(\tau')\mathrm{d}\tau'}. \tag{S10}$$

Assume that $\Delta_{\mathrm{tQ}}(\tau')$ changes rather slowly over $\tau'$ to satisfy

$$\Delta_{\mathrm{tQ}}(\tau') \approx \Delta_{\mathrm{tQ}}(\tau) \text{ for } \tau' \text{ in the range } \tau - \Delta_{\mathrm{tQ}}^{-1}(\tau) \lesssim \tau' \le \tau. \tag{S11}$$

In physiologically-relevant conditions, Eq. (S11) is satisfied as readily as Eq. (S7) (Text S5). With Eq. (S11), Eq. (S10) for $\tau \gg \tau_0 + \Delta_{\mathrm{tQ}}^{-1}(\tau_0)$ is further approximated as

$$\bar{C}(\tau) \approx \Delta_{\mathrm{tQ}}(\tau) \int_{-\infty}^{\tau} \bar{C}_{\mathrm{tQ}}(\tau') e^{-(\tau-\tau')\Delta_{\mathrm{tQ}}(\tau)} \mathrm{d}\tau'. \tag{S12}$$

The Taylor expansion $\bar{C}_{\mathrm{tQ}}(\tau') = \bar{C}_{\mathrm{tQ}}(\tau) - (\tau-\tau')\bar{C}'_{\mathrm{tQ}}(\tau) + (\tau-\tau')^2 \bar{C}''_{\mathrm{tQ}}(\tau)/2 - \cdots$ leads Eq. (S12) to

$$\begin{aligned}\bar{C}(\tau) &\approx \bar{C}_{\mathrm{tQ}}(\tau) - \Delta_{\mathrm{tQ}}^{-1}(\tau)\frac{\mathrm{d}\bar{C}_{\mathrm{tQ}}(\tau)}{\mathrm{d}\tau}\int_0^\infty x e^{-x}\mathrm{d}x + \frac{\Delta_{\mathrm{tQ}}^{-2}(\tau)}{2}\frac{\mathrm{d}^2\bar{C}_{\mathrm{tQ}}(\tau)}{\mathrm{d}\tau^2}\int_0^\infty x^2 e^{-x}\mathrm{d}x - \cdots \\ &= \bar{C}_{\mathrm{tQ}}(\tau) - \Delta_{\mathrm{tQ}}^{-1}(\tau)\frac{\mathrm{d}\bar{C}_{\mathrm{tQ}}(\tau)}{\mathrm{d}\tau} + \Delta_{\mathrm{tQ}}^{-2}(\tau)\frac{\mathrm{d}^2\bar{C}_{\mathrm{tQ}}(\tau)}{\mathrm{d}\tau^2} - \cdots \\ &= \bar{C}_{\mathrm{tQe}}(\tau) + \Delta_{\mathrm{tQ}}^{-2}(\tau)\frac{\mathrm{d}^2\bar{C}_{\mathrm{tQ}}(\tau)}{\mathrm{d}\tau^2} - \cdots,\end{aligned} \tag{S13}$$

where $\bar{C}_{\mathrm{tQe}}(\tau)$ is defined as

$$\bar{C}_{\mathrm{tQe}}(\tau) \equiv \bar{C}_{\mathrm{tQ}}(\tau) - \Delta_{\mathrm{tQ}}^{-1}(\tau)\frac{\mathrm{d}\bar{C}_{\mathrm{tQ}}(\tau)}{\mathrm{d}\tau}. \tag{S14}$$

For the approximation of $\bar{C}(\tau)$, one may be tempted to use $\bar{C}_{\mathrm{tQe}}(\tau)$ in Eq. (S14). However, as proven in Text S2, the sheer use of $\bar{C}_{\mathrm{tQe}}(\tau)$ is susceptible to the overestimation of the amplitude of $\bar{C}(\tau)$ when $\bar{C}(\tau)$ is rhythmic over time. To detour this overestimation problem, we take the Taylor expansion of the time-delayed form of $\bar{C}_{\mathrm{tQ}}(\tau)$:

$$\bar{C}_{\mathrm{tQ}}\left[\tau - \Delta_{\mathrm{tQ}}^{-1}(\tau)\right] = \bar{C}_{\mathrm{tQ}}(\tau) - \Delta_{\mathrm{tQ}}^{-1}(\tau)\frac{\mathrm{d}\bar{C}_{\mathrm{tQ}}(\tau)}{\mathrm{d}\tau} + \frac{\Delta_{\mathrm{tQ}}^{-2}(\tau)}{2}\frac{\mathrm{d}^2\bar{C}_{\mathrm{tQ}}(\tau)}{\mathrm{d}\tau^2} - \cdots. \tag{S15}$$

Strikingly, the zeroth-order and first-order derivative terms of $\bar{C}_{\mathrm{tQ}}(\tau)$ on the right-hand side above are identical to $\bar{C}_{\mathrm{tQe}}(\tau)$, and the second-order derivative term still covers a half of that term in Eq. (S13). Hence, $\bar{C}_{\mathrm{tQ}}\left[\tau - \Delta_{\mathrm{tQ}}^{-1}(\tau)\right]$ bears the potential for the approximants of $\bar{C}(\tau)$. Besides, the overestimation of the amplitude of rhythmic $\bar{C}(\tau)$ by $\bar{C}_{\mathrm{tQ}}\left[\tau - \Delta_{\mathrm{tQ}}^{-1}(\tau)\right]$ would not be as serious as by $\bar{C}_{\mathrm{tQe}}(\tau)$ and at worst equals that by $\bar{C}_{\mathrm{tQ}}(\tau)$, because $\bar{C}_{\mathrm{tQ}}\left[\tau - \Delta_{\mathrm{tQ}}^{-1}(\tau)\right]$ and $\bar{C}_{\mathrm{tQ}}(\tau)$ themselves have the same amplitudes.

One caveat with the use of $\bar{C}_{\mathrm{tQ}}\left[\tau - \Delta_{\mathrm{tQ}}^{-1}(\tau)\right]$ to estimate $\bar{C}(\tau)$ is that $\bar{C}_{\mathrm{tQ}}\left[\tau - \Delta_{\mathrm{tQ}}^{-1}(\tau)\right]$ may not necessarily satisfy the relation $\bar{C}_{\mathrm{tQ}}\left[\tau - \Delta_{\mathrm{tQ}}^{-1}(\tau)\right] \leq \min\left[\overline{A}(\tau), \overline{B}(\tau)\right]$ favored by Eq. (S2). As a practical safeguard to avoid this problem, we propose the following approximant for $\bar{C}(\tau)$ consistent with Eq. (S2):

$$\bar{C}_{\gamma}(\tau) \equiv \min\left\{\bar{C}_{\mathrm{tQ}}\left[\tau - \Delta_{\mathrm{tQ}}^{-1}(\tau)\right], \bar{A}(\tau), \bar{B}(\tau)\right\}. \tag{S16}$$

We refer to this formulation as the ETS, and its correspondent for $C(t)$ is $C_\gamma(t)$ in Eq. (6). The delay term $\Delta_{\mathrm{tQ}}^{-1}(\tau)$ in Eq. (S16), that is $\left[k_\delta \Delta_{\mathrm{tQ}}(t)\right]^{-1}$ in the domain of time $t$, is interpreted as the molecular relaxation time in complex formation: this interpretation comes from Eq. (S12) where the memory of instantaneous A and B concentrations in $\bar{C}_{\mathrm{tQ}}(\tau')$ has the duration with the time-scale of $\Delta_{\mathrm{tQ}}^{-1}(\tau)$ during the formation of complex AB. To identify the underlying factors of this relaxation time, we rewrite Eq. (S8) as

$$\frac{\mathrm{d}}{\mathrm{d}\tau}\left[\bar{C}(\tau) - \bar{C}_{\mathrm{tQ}}(\tau)\right] \approx -\frac{\mathrm{d}\bar{C}_{\mathrm{tQ}}(\tau)}{\mathrm{d}\tau} - \left[\Delta_{\mathrm{tQ}}(\tau) - 1\right]\left[\bar{C}(\tau) - \bar{C}_{\mathrm{tQ}}(\tau)\right] - \left[\bar{C}(\tau) - \bar{C}_{\mathrm{tQ}}(\tau)\right]. \quad \text{(S17)}$$

Regarding the above relaxation dynamics, notice that $\Delta_{\mathrm{tQ}}(\tau) - 1 = \overline{A}(\tau) + \overline{B}(\tau) - 2\bar{C}_{\mathrm{tQ}}(\tau) = \left[\overline{A}(\tau) - \bar{C}_{\mathrm{tQ}}(\tau)\right] + \left[\overline{B}(\tau) - \bar{C}_{\mathrm{tQ}}(\tau)\right] \geq 0$ from Eq. (S4). In other words, $\Delta_{\mathrm{tQ}}(\tau) - 1$ indicates the total free molecule abundanceat the quasi-steady state. Furthermore, the second and third terms on the right-hand side of Eq. (S17) are respectively contributed to by $[\bar{A}(\tau) - \bar{C}(\tau)][\bar{B}(\tau) - \bar{C}(\tau)]$ and $-\bar{C}(\tau)$ on the right-hand side of Eq. (S1). Because $[\bar{A}(\tau) - \bar{C}(\tau)][\bar{B}(\tau) - \bar{C}(\tau)]$ is a free molecule binding rate, the second term on the right-hand side of Eq. (S17) reflects a decrease in the free molecule binding rate with an increase in the complex abundance that depletes the free molecules. This effect results in the shorter relaxation time than expected only by the decay of the complex reflected in the third term. Because the second term is proportional to $\Delta_{\mathrm{tQ}}(\tau) - 1$, the free molecule depletion effect increases with the free molecule availability. Therefore, the relaxation time takes a decreasing function of the free molecule availability, as the form of the delay term $\Delta_{\mathrm{tQ}}^{-1}(\tau)$ in Eq. (S16).

Related to the relaxation time, Eq. (S16) is ill-defined for $\tau - \Delta_{\mathrm{tQ}}^{-1}(\tau) < \tau_0$ where $\tau_0$ is an initial point of $\tau$. In fact, from the interpretation of $\Delta_{\mathrm{tQ}}^{-1}(\tau)$ as the relaxation time, $\tau$ should satisfy $\tau \gg \tau_0 + \Delta_{\mathrm{tQ}}^{-1}(\tau_0)$ for any rate law (e.g., the ETS, tQSSA, or sQSSA) whose form does not depend on the initial conditions. This point is also evident from the last term in Eq. (S10).

**Text S2. Amplitude overestimation with Eq. (S14)**

We here consider a situation that $\bar{C}(\tau)$ in Eq. (S3) is rhythmic over time. At the peak or trough time of $\bar{C}(\tau)$, $\bar{C}'(\tau) = 0$ and therefore $\bar{C}(\tau) = \bar{C}_{\mathrm{tQ}}(\tau)$ at that time. Combined with $\min_\tau\left[\bar{C}_{\mathrm{tQ}}(\tau)\right] \leq \bar{C}_{\mathrm{tQ}}(\tau) \leq \max_\tau\left[\bar{C}_{\mathrm{tQ}}(\tau)\right]$, it leads to $\max_\tau[\bar{C}(\tau)] \leq \max_\tau\left[\bar{C}_{\mathrm{tQ}}(\tau)\right]$ and $\min_\tau[\bar{C}(\tau)] \geq \min_\tau\left[\bar{C}_{\mathrm{tQ}}(\tau)\right]$.

In addition, at the peak or trough time of $\bar{C}_{\mathrm{tQ}}(\tau)$, $\bar{C}'_{\mathrm{tQ}}(\tau) = 0$ in Eq. (S14) and therefore $\bar{C}_{\mathrm{tQ}}(\tau) = \bar{C}_{\mathrm{tQe}}(\tau)$ at that time. It similarly leads to $\max_\tau[\bar{C}_{\mathrm{tQ}}(\tau)] \le \max_\tau[\bar{C}_{\mathrm{tQe}}(\tau)]$ and $\min_\tau[\bar{C}_{\mathrm{tQ}}(\tau)] \ge \min_\tau[\bar{C}_{\mathrm{tQe}}(\tau)]$.

Taken together, $\max_\tau[\bar{C}(\tau)] \le \max_\tau[\bar{C}_{\mathrm{tQ}}(\tau)] \le \max_\tau[\bar{C}_{\mathrm{tQe}}(\tau)]$ and $\min_\tau[\bar{C}(\tau)] \ge \min_\tau[\bar{C}_{\mathrm{tQ}}(\tau)] \ge \min_\tau[\bar{C}_{\mathrm{tQe}}(\tau)]$. As a result, $\bar{C}_{\mathrm{tQ}}(\tau)$ tends to overestimate the amplitude of $\bar{C}(\tau)$ but $\bar{C}_{\mathrm{tQe}}(\tau)$ does more seriously, through the over- and under-estimation of the peak and trough levels of $\bar{C}(\tau)$, respectively.

**Text S3. Amplitude overestimation with simpler new rate laws**

We here show that, unlike the ETS, any simpler new rate law without a time-delay term would not properly work for actively time-varying molecular complex levels, because this type of a rate law is susceptible to amplitude overestimation.

As a new rate law without a time-delay term, one may suggest a certain function of $A(t)$, $B(t)$, $A'(t)$, and $B'(t)$. Here, $A(t)$ and $B(t)$ are the concentrations of A and B molecules in Eq. (1), and $A'(t)$ and $B'(t)$ are their time derivatives to reflect the past trajectory in place of a time-delay term. Further consideration of higher-order derivatives is likely to make the functional form even more complicated than with only the time-delay term, and we thus discard the higher-order terms.

For simplicity, suppose that $A(t)$ is rhythmic over time while $B(t)$ is constant. In this case, the above function just takes the form of $f[A(t), A'(t)]$. This function should satisfy $f[A(t), 0] = C_{\mathrm{tQ}}(t)$ with $C_{\mathrm{tQ}}(t)$ in Eq. (2) as the exact steady-state solution of Eq. (1). At the peak and trough times of $A(t)$ [i.e., $A'(t) = 0$], $C_{\mathrm{tQ}}(t)$ in its definition reaches $\max_t[C_{\mathrm{tQ}}(t)]$ and $\min_t[C_{\mathrm{tQ}}(t)]$, respectively. Together with this fact, $\min_t\{f[A(t), A'(t)]\} \le f[A(t), A'(t)]|_{A'(t)=0} \le \max_t\{f[A(t), A'(t)]\}$ and the above $f[A(t), 0] = C_{\mathrm{tQ}}(t)$ lead to $\max_t[C_{\mathrm{tQ}}(t)] \le \max_t\{f[A(t), A'(t)]\}$ and $\min_t[C_{\mathrm{tQ}}(t)] \ge \min_t\{f[A(t), A'(t)]\}$.

In addition, at the peak or trough time of $C(t)$ in Eq. (1), $C'(t) = 0$ and therefore $C(t) = C_{\mathrm{tQ}}(t)$ at that time. Combined with $\min_t[C_{\mathrm{tQ}}(t)] \le C_{\mathrm{tQ}}(t) \le \max_t[C_{\mathrm{tQ}}(t)]$, it leads to $\max_t[C(t)] \le \max_t[C_{\mathrm{tQ}}(t)]$ and $\min_t[C(t)] \ge \min_t[C_{\mathrm{tQ}}(t)]$.

Taken together, $\max_t[C(t)] \le \max_t[C_{\mathrm{tQ}}(t)] \le \max_t\{f[A(t), A'(t)]\}$ and $\min_t[C(t)] \ge \min_t[C_{\mathrm{tQ}}(t)] \ge \min_t\{f[A(t), A'(t)]\}$. As a result, $C_{\mathrm{tQ}}(t)$ tends to overestimate the amplitude of $C(t)$ but $f[A(t), A'(t)]$ does more seriously, through the over- and under-estimation of the peak and trough levels of $C(t)$, respectively. Therefore, $f[A(t), A'(t)]$ as a simple rate law without a time-delay term is prone to the amplitude overestimation.

**Text S4. Rate law derivation for TF–DNA interactions**

Imagine that a TF binds to a DNA molecule in the nucleus. The highly discrete nature of the TF–DNA binding number does not allow the use of Eq. (1) that involves the time derivative of $C(t)$ with its assumed continuity. Instead of Eq. (1), we can use the chemical master equation [S10] to rigorously describe the TF–DNA binding dynamics. Let $P(n,t)$ denote the probability that $n$ copies of the TF are occupying the DNA site at time $t$. If this DNA site can afford at most $N$ copies of the TF at once, $n = 0, 1, \cdots, N$ and $\sum_{n=0}^{N} P(n,t) = 1$. If we further define $P(n,t) \equiv 0$ for $n \neq 0, 1, \cdots, N$ and assume that the DNA-binding TFs are hardly accessible by molecular machineries such as for protein degradation, the temporal change of $P(n,t)$ with $n = 0, 1, \cdots, N$ is governed by the following master equation:

$$\frac{\partial P(n,t)}{\partial t} = k_{\mathrm{a}} V \left[A_{\mathrm{TF}}(t) - \frac{n-1}{V}\right]\left(B_{\mathrm{DNA}} - \frac{n-1}{V}\right) P(n-1,t) - \left\{k_{\mathrm{a}} V \left[A_{\mathrm{TF}}(t) - \frac{n}{V}\right]\left(B_{\mathrm{DNA}} - \frac{n}{V}\right) + n k_{\delta}\right\} P(n,t) + (n+1) k_{\delta} P(n+1,t), \tag{S18}$$

where $k_{\mathrm{a}}$ and $k_{\delta}$ denote the TF–DNA binding and unbinding rates, respectively, $V$ is the nuclear volume, $A_{\mathrm{TF}}(t)$ is the total TF concentration in the nucleus, and $B_{\mathrm{DNA}}$ is the "concentration" of the target DNA site, i.e., $B_{\mathrm{DNA}} = NV^{-1}$. Here, we assume $A_{\mathrm{TF}}(t)$ to be uniquely determined at each time $t$ with little stochasticity in $A_{\mathrm{TF}}(t)$ itself and a steady nuclear volume with constant $V$.

Introducing a quantity $C_{\mathrm{TF}}(t) \equiv \langle nV^{-1} \rangle = V^{-1} \sum_{n=0}^{N} nP(n,t)$ to Eq. (S18) results in

$$\begin{aligned}\frac{\mathrm{d}C_{\mathrm{TF}}(t)}{\mathrm{d}t} &= k_{\mathrm{a}} \sum_{n=0}^{N} \left[A_{\mathrm{TF}}(t) - \frac{n}{V}\right]\left(B_{\mathrm{DNA}} - \frac{n}{V}\right) P(n,t) - k_{\delta} C_{\mathrm{TF}}(t) \\ &= k_{\mathrm{a}} \left\{[A_{\mathrm{TF}}(t) - C_{\mathrm{TF}}(t)][B_{\mathrm{DNA}} - C_{\mathrm{TF}}(t)] + \langle \left(\tfrac{n}{V}\right)^2 \rangle - \langle \tfrac{n}{V} \rangle^2 \right\} - k_{\delta} C_{\mathrm{TF}}(t),\end{aligned} \tag{S19}$$

where $\langle (nV^{-1})^2 \rangle = V^{-2} \sum_{n=0}^{N} n^2 P(n,t)$. Eq. (S19) is reminiscent of Eq. (1), when the stochastic fluctuation in the TF binding $[\langle (nV^{-1})^2 \rangle - \langle nV^{-1} \rangle^2]$ is negligible. The stochastic fluctuation, however, cannot be ignored for small $N$. For simplicity, we will henceforth consider the case of $N = 1$ and thus of $B_{\mathrm{DNA}} = V^{-1}$. In this case, Eq. (S19) is rewritten as

$$\frac{\mathrm{d}\bar{C}_{\mathrm{TF}}(\tau)}{\mathrm{d}\tau} = \frac{\bar{A}_{\mathrm{TF}}(\tau)}{KV} - [1 + \bar{A}_{\mathrm{TF}}(\tau)]\bar{C}_{\mathrm{TF}}(\tau) = [1 + \bar{A}_{\mathrm{TF}}(\tau)]\left[\bar{C}_{\mathrm{TFQ}}(\tau) - \bar{C}_{\mathrm{TF}}(\tau)\right], \tag{S20}$$

where $\tau \equiv k_{\delta} t$, $K \equiv k_{\delta}/k_{\mathrm{a}}$, $\bar{C}_{\mathrm{TF}}(\tau) \equiv C_{\mathrm{TF}}(t)/K$, $\bar{A}_{\mathrm{TF}}(\tau) \equiv A_{\mathrm{TF}}(t)/K$, and

$$\bar{C}_{\mathrm{TFQ}}(\tau) \equiv \frac{\bar{A}_{\mathrm{TF}}(\tau)}{KV[1+\bar{A}_{\mathrm{TF}}(\tau)]}. \tag{S21}$$

Eq. (S21) is the dimensionless form of Eq. (7) with $\bar{C}_{\mathrm{TFQ}}(\tau) = C_{\mathrm{TFQ}}(t)/K$.

Here, the quasi-steady state assumption is that $\bar{C}_{\mathrm{TF}}(\tau)$ approaches an equilibrium (quasi-steady state) fast enough each time, given the value of $\bar{A}_{\mathrm{TF}}(\tau)$. Because $\bar{C}'_{\mathrm{TF}}(\tau) \to 0$ in Eq. (S20) when $\bar{C}_{\mathrm{TF}}(\tau) \to \bar{C}_{\mathrm{TFQ}}(\tau)$ given the value of $\bar{A}_{\mathrm{TF}}(\tau)$, we take the approximation $\bar{C}_{\mathrm{TF}}(\tau) \approx \bar{C}_{\mathrm{TFQ}}(\tau)$, or equivalently, $C_{\mathrm{TF}}(t) \approx C_{\mathrm{TFQ}}(t)$ under the quasi-steady state assumption. As discussed after Eq. (7), this approximation is neither exactly the tQSSA nor sQSSA, and we thus refer to it as the QSSA for the TF–DNA interactions. Of note, $\bar{C}_{\mathrm{TFQ}}(\tau)$ corresponds to a special case of the previously-studied, stochastic QSSA [S11,S12] for arbitrary molecular copy numbers such as for multiple DNA binding sites.

To improve the rate law for time-varying TF concentration beyond the quasi-steady state assumption, notice that the exact solution of Eq. (S20) is

$$\bar{C}_{\mathrm{TF}}(\tau) = \int_{\tau_0}^{\tau} [1 + \bar{A}_{\mathrm{TF}}(\tau')]\bar{C}_{\mathrm{TFQ}}(\tau') e^{-\int_{\tau'}^{\tau}[1+\bar{A}_{\mathrm{TF}}(\tau'')]\mathrm{d}\tau''} \mathrm{d}\tau' + \bar{C}_{\mathrm{TF}}(\tau_0) e^{-\int_{\tau_0}^{\tau}[1+\bar{A}_{\mathrm{TF}}(\tau')]\mathrm{d}\tau'}, \tag{S22}$$

where $\tau_0$ denotes an arbitrarily assigned, initial point of $\tau$. Assume that $1 + \bar{A}_{\mathrm{TF}}(\tau')$ changes rather slowly over $\tau'$ to satisfy

$$1 + \bar{A}_{\mathrm{TF}}(\tau') \approx 1 + \bar{A}_{\mathrm{TF}}(\tau) \text{ for } \tau' \text{ in the range } \tau - \frac{1}{1+\bar{A}_{\mathrm{TF}}(\tau)} \lesssim \tau' \leq \tau. \tag{S23}$$

In physiologically-relevant conditions, Eq. (S23) is readily satisfied (see Text S5). With Eq. (S23), Eq. (S22) for $\tau \gg \tau_0 + [1 + \bar{A}_{\mathrm{TF}}(\tau_0)]^{-1}$ is approximated as

$$\bar{C}_{\mathrm{TF}}(\tau) \approx [1 + \bar{A}_{\mathrm{TF}}(\tau)] \int_{-\infty}^{\tau} \bar{C}_{\mathrm{TFQ}}(\tau') e^{-[1+\bar{A}_{\mathrm{TF}}(\tau)](\tau-\tau')} \mathrm{d}\tau'. \tag{S24}$$

The Taylor expansion $\bar{C}_{\mathrm{TFQ}}(\tau') = \bar{C}_{\mathrm{TFQ}}(\tau) - (\tau - \tau')\bar{C}'_{\mathrm{TFQ}}(\tau) + (\tau - \tau')^2 \bar{C}''_{\mathrm{TFQ}}(\tau)/2 - \cdots$ leads Eq. (S24) to

$$\begin{aligned}
\bar{C}_{\mathrm{TF}}(\tau) &\approx \bar{C}_{\mathrm{TFQ}}(\tau) - \frac{1}{1 + \bar{A}_{\mathrm{TF}}(\tau)} \frac{\mathrm{d}\bar{C}_{\mathrm{TFQ}}(\tau)}{\mathrm{d}\tau} \int_0^{\infty} x e^{-x} \mathrm{d}x \\
&\quad + \frac{1}{2[1 + \bar{A}_{\mathrm{TF}}(\tau)]^2} \frac{\mathrm{d}^2\bar{C}_{\mathrm{TFQ}}(\tau)}{\mathrm{d}\tau^2} \int_0^{\infty} x^2 e^{-x} \mathrm{d}x - \cdots \\
&= \bar{C}_{\mathrm{TFQ}}(\tau) - \frac{1}{1+\bar{A}_{\mathrm{TF}}(\tau)} \frac{\mathrm{d}\bar{C}_{\mathrm{TFQ}}(\tau)}{\mathrm{d}\tau} + \frac{1}{[1+\bar{A}_{\mathrm{TF}}(\tau)]^2} \frac{\mathrm{d}^2\bar{C}_{\mathrm{TFQ}}(\tau)}{\mathrm{d}\tau^2} - \cdots \\
&= \bar{C}_{\mathrm{TFQe}}(\tau) + \frac{1}{[1+\bar{A}_{\mathrm{TF}}(\tau)]^2} \frac{\mathrm{d}^2\bar{C}_{\mathrm{TFQ}}(\tau)}{\mathrm{d}\tau^2} - \cdots,
\end{aligned} \tag{S25}$$

where $\bar{C}_{\mathrm{TFQe}}(\tau)$ is defined as

$$\bar{C}_{\mathrm{TFQe}}(\tau) \equiv \bar{C}_{\mathrm{TFQ}}(\tau) - \frac{1}{1+\bar{A}_{\mathrm{TF}}(\tau)} \frac{\mathrm{d}\bar{C}_{\mathrm{TFQ}}(\tau)}{\mathrm{d}\tau}. \tag{S26}$$

On the other hand, the Taylor expansion of the time-delayed form of $\bar{C}_{\mathrm{TFQ}}(\tau)$ is

$$\bar{C}_{\mathrm{TFQ}}\left[\tau - \frac{1}{1+\bar{A}_{\mathrm{TF}}(\tau)}\right] = \bar{C}_{\mathrm{TFQ}}(\tau) - \frac{1}{1+\bar{A}_{\mathrm{TF}}(\tau)} \frac{\mathrm{d}\bar{C}_{\mathrm{TFQ}}(\tau)}{\mathrm{d}\tau} + \frac{1}{2[1+\bar{A}_{\mathrm{TF}}(\tau)]^2} \frac{\mathrm{d}^2\bar{C}_{\mathrm{TFQ}}(\tau)}{\mathrm{d}\tau^2} - \cdots. \tag{S27}$$

Interestingly, the zeroth-order and first-order derivative terms of $\bar{C}_{\mathrm{TFQ}}(\tau)$ on the right-hand side above are identical to $\bar{C}_{\mathrm{TFQe}}(\tau)$, and the second-order derivative term still covers a half of that term in Eq. (S25). Hence, we propose the following approximant for $\bar{C}_{\mathrm{TF}}(\tau)$:

$$\bar{C}_{\mathrm{TF}\gamma}(\tau) \equiv \bar{C}_{\mathrm{TFQ}}\left[\tau - \frac{1}{1+\bar{A}_{\mathrm{TF}}(\tau)}\right]. \tag{S28}$$

This formulation is the ETS of the TF–DNA interaction. The corresponding approximant for $C_{\mathrm{TF}}(t)$ is $C_{\mathrm{TF}\gamma}(t)$ in Eq. (8). Of note, Eq. (S28) is ill-defined for $\tau - [1 + \bar{A}_{\mathrm{TF}}(\tau)]^{-1} < \tau_0$ where $\tau_0$ is an initial point of $\tau$. In fact, from the last term in Eq. (S22), $\tau$ should satisfy $\tau \gg \tau_0 + [1 + \bar{A}_{\mathrm{TF}}(\tau_0)]^{-1}$ for the application of any rate law (e.g., the ETS or QSSA) whose form does not depend on the initial conditions.

**Text S5. Preconditions of rate laws**

We here clarify the preconditions of the ETS as well as those of the (t)QSSA. To first show the preconditions of the ETS with $C_\gamma(t)$ in Eq. (6), or equivalently $\bar{C}_\gamma(\tau)$ in Eq. (S16), we revisit the condition in Eq. (S7). Replacing $\bar{C}(\tau)$ in Eq. (S7) by $\bar{C}_{\mathrm{tQe}}(\tau)$ in Eq. (S14) leads to the following self-consistency condition:

$$\varepsilon_1(\tau) \equiv \Delta_{\mathrm{tQ}}^{-2}(\tau) \left|\frac{\mathrm{d}\bar{C}_{\mathrm{tQ}}(\tau)}{\mathrm{d}\tau}\right| \ll 1, \tag{S29}$$

where $\bar{C}'_{\mathrm{tQ}}(\tau)$ is given by

$$\frac{\mathrm{d}\bar{C}_{\mathrm{tQ}}(\tau)}{\mathrm{d}\tau} = \Delta_{\mathrm{tQ}}^{-1}(\tau)\{[\bar{B}(\tau) - \bar{C}_{\mathrm{tQ}}(\tau)]\bar{A}(\tau)\mu_{\mathrm{A}}(\tau) + [\bar{A}(\tau) - \bar{C}_{\mathrm{tQ}}(\tau)]\bar{B}(\tau)\mu_{\mathrm{B}}(\tau)\}. \tag{S30}$$

Here, $\mu_{\mathrm{A}}(\tau) \equiv \bar{A}'(\tau)/\bar{A}(\tau)$ , $\mu_{\mathrm{B}}(\tau) \equiv \bar{B}'(\tau)/\bar{B}(\tau)$, and the derivation of Eq. (S30) is straightforward from Eq. (S4). For the sake of simplicity, we will keep using these notations $\mu_{\mathrm{A}}(\tau)$ and $\mu_{\mathrm{B}}(\tau)$. Next, we revisit another condition in Eq. (S11). By applying an expansion $\Delta_{\mathrm{tQ}}(\tau') \approx \Delta_{\mathrm{tQ}}(\tau) + (\tau' - \tau)\Delta'_{\mathrm{tQ}}(\tau)$ to Eq. (S11), we obtain $|(\tau' - \tau)\Delta'_{\mathrm{tQ}}(\tau)| \ll \Delta_{\mathrm{tQ}}(\tau)$ for $\tau'$

in the range $\tau - \Delta_{\rm tQ}^{-1}(\tau) \lesssim \tau' \leq \tau$ and therefore $\Delta_{\rm tQ}^{-2}(\tau)\left|\Delta_{\rm tQ}'(\tau)\right| \ll 1$. By the definition of $\Delta_{\rm tQ}(\tau)$ in Eq. (S5), this condition is equivalent to

$$\varepsilon_2(\tau) \equiv \Delta_{\rm tQ}^{-3}(\tau)|[1+\bar{A}(\tau)-\bar{B}(\tau)]\bar{A}(\tau)\mu_{\rm A}(\tau)+[1+\bar{B}(\tau)-\bar{A}(\tau)]\bar{B}(\tau)\mu_{\rm B}(\tau)| \ll 1. \quad \text{(S31)}$$

The last condition below arises from the comparison between Eqs. (S13) and (S15), which show the difference of $\sim\Delta_{\rm tQ}^{-2}(\tau)\bar{C}_{\rm tQ}''(\tau)/2$:

$$\varepsilon_\gamma(\tau) \equiv \frac{\Delta_{\rm tQ}^{-2}(\tau)}{2\bar{C}_{\rm tQ}[\tau-\Delta_{\rm tQ}^{-1}(\tau)]}\left|\frac{{\rm d}^2\bar{C}_{\rm tQ}(\tau)}{{\rm d}\tau^2}\right| \ll 1, \quad \text{(S32)}$$

where $\bar{C}_{\rm tQ}''(\tau)$ is obtained from Eq. (S30) as

$$\frac{{\rm d}^2\bar{C}_{\rm tQ}(\tau)}{{\rm d}\tau^2} = \Delta_{\rm tQ}^{-3}(\tau)\Big\{\big\{\Delta_{\rm tQ}^2(\tau)-2\bar{A}(\tau)\big[1+\bar{A}(\tau)-\bar{C}_{\rm tQ}(\tau)\big]\big\}\big[\bar{B}(\tau)-\bar{C}_{\rm tQ}(\tau)\big]\bar{A}(\tau)\mu_{\rm A}^2(\tau)+ \big\{\Delta_{\rm tQ}^2(\tau)-2\bar{B}(\tau)\big[1+\bar{B}(\tau)-\bar{C}_{\rm tQ}(\tau)\big]\big\}\big[\bar{A}(\tau)-\bar{C}_{\rm tQ}(\tau)\big]\bar{B}(\tau)\mu_{\rm B}^2(\tau)+\big\{1+\Delta_{\rm tQ}^2(\tau)- [\bar{A}(\tau)-\bar{B}(\tau)]^2\big\}\bar{A}(\tau)\bar{B}(\tau)\mu_{\rm A}(\tau)\mu_{\rm B}(\tau)+\Delta_{\rm tQ}^2(\tau)\Big\{\big[\bar{B}(\tau)-\bar{C}_{\rm tQ}(\tau)\big]\bar{A}(\tau)\frac{{\rm d}\mu_{\rm A}(\tau)}{{\rm d}\tau}+ \big[\bar{A}(\tau)-\bar{C}_{\rm tQ}(\tau)\big]\bar{B}(\tau)\frac{{\rm d}\mu_{\rm B}(\tau)}{{\rm d}\tau}\Big\}\Big\}. \quad \text{(S33)}$$

In summary, our approximant $\bar{C}_\gamma(\tau)$ shall work when Eqs. (S29), (S31), and (S32) are satisfied. As we will show later, Eqs. (S29) and (S31) are in fact easy to satisfy and thus only Eq. (S32) tends to serve as the relevant factor of the validity of $\bar{C}_\gamma(\tau)$.

On the other hand, the tQSSA in Eq. (2) or (S4) would be valid in the following condition from Eq. (S13), instead of the condition in Eq. (S32):

$$\varepsilon_{\rm tQ}(\tau) \equiv \frac{1}{\bar{C}_{\rm tQ}(\tau)}\left|\Delta_{\rm tQ}^{-1}(\tau)\frac{{\rm d}\bar{C}_{\rm tQ}(\tau)}{{\rm d}\tau}-\Delta_{\rm tQ}^{-2}(\tau)\frac{{\rm d}^2\bar{C}_{\rm tQ}(\tau)}{{\rm d}\tau^2}\right| \ll 1, \quad \text{(S34)}$$

where $\bar{C}_{\rm tQ}'(\tau)$ and $\bar{C}_{\rm tQ}''(\tau)$ are given by Eqs. (S30) and (S33), respectively. Note that this condition for the tQSSA is more rigorous than the previously-reported condition [S3]. If molecular concentrations vary over time with a characteristic time-scale of $T$, Eq. (S34) roughly requires the effective time delay $\left[k_\delta\Delta_{\rm tQ}(t)\right]^{-1}$ in Eq. (6) to be smaller enough than $T$ for the validity of the tQSSA.

Regarding $\bar{C}(\tau)$ from Eqs. (S1) and (S40) in Text S7, one may expect that the high accuracy of the ETS compared to the tQSSA might be indicated by the range of its valid conditions in Eqs. (S29), (S31), and (S32). In fact, most of the physiologically-relevant conditions in Table S3 (88.1%) satisfy both $\max_\tau[\varepsilon_1(\tau)] \leq 0.1$ and $\max_\tau[\varepsilon_2(\tau)] \leq 0.1$, and therefore only $\varepsilon_\gamma(\tau)$ and $\varepsilon_{\rm tQ}(\tau)$ remain the key determinants of the validities of the ETS and tQSSA, respectively. Our analysis reveals that $\max_\tau\left[\varepsilon_\gamma(\tau)\right] \leq 0.1$ for 55.9% of the simulated conditions and

$\max_\tau[\varepsilon_{\mathrm{tQ}}(\tau)] \le 0.1$ for 45.4% of the same conditions. Among them, the conditions with $\max_\tau[\varepsilon_\gamma(\tau)] \le 0.1$ cover 99.8% of the conditions with $\max_\tau[\varepsilon_{\mathrm{tQ}}(\tau)] \le 0.1$, supporting more general applicability of the ETS than the tQSSA's. In addition, Fig. S1(a) shows that the ETS is particularly more valid than the tQSSA for larger $K$ and smaller $k_\delta$, which tend to lengthen the effective time delay $[k_\delta \Delta_{\mathrm{tQ}}(t)]^{-1}$.

Next, to clarify the preconditions of the ETS with $C_{\mathrm{TF}\gamma}(t)$ in Eq. (8), or equivalently $\bar{C}_{\mathrm{TF}\gamma}(\tau)$ in Eq. (S28), we first revisit the condition in Eq. (S23). Applying an expansion $1 + \bar{A}_{\mathrm{TF}}(\tau') \approx 1 + \bar{A}_{\mathrm{TF}}(\tau) + (\tau' - \tau)\bar{A}'_{\mathrm{TF}}(\tau)$ to Eq. (S23) gives rise to this condition:

$$\varepsilon_{\mathrm{TF}}(\tau) \equiv \frac{\bar{A}_{\mathrm{TF}}(\tau)}{[1+\bar{A}_{\mathrm{TF}}(\tau)]^2} |\mu_{\mathrm{TF}}(\tau)| \ll 1, \tag{S35}$$

where $\mu_{\mathrm{TF}}(\tau) \equiv \bar{A}'_{\mathrm{TF}}(\tau)/\bar{A}_{\mathrm{TF}}(\tau)$. In addition, the comparison between Eqs. (S25) and (S27) shows the difference of $\sim\bar{C}''_{\mathrm{TFQ}}(\tau)/\{2[1 + \bar{A}_{\mathrm{TF}}(\tau)]^2\}$ and thereby offers this condition:

$$\varepsilon_{\mathrm{TF}\gamma}(\tau) \equiv \frac{1}{2[1+\bar{A}_{\mathrm{TF}}(\tau)]^2 \bar{C}_{\mathrm{TFQ}}\{\tau-[1+\bar{A}_{\mathrm{TF}}(\tau)]^{-1}\}} \left|\frac{\mathrm{d}^2\bar{C}_{\mathrm{TFQ}}(\tau)}{\mathrm{d}\tau^2}\right| \ll 1, \tag{S36}$$

where $\bar{C}''_{\mathrm{TFQ}}(\tau)$ is obtained from Eq. (S21) as

$$\frac{\mathrm{d}^2\bar{C}_{\mathrm{TFQ}}(\tau)}{\mathrm{d}\tau^2} = \frac{\bar{C}_{\mathrm{TFQ}}(\tau)}{1+\bar{A}_{\mathrm{TF}}(\tau)} \left[\frac{1-\bar{A}_{\mathrm{TF}}(\tau)}{1+\bar{A}_{\mathrm{TF}}(\tau)} \mu^2_{\mathrm{TF}}(\tau) + \frac{\mathrm{d}\mu_{\mathrm{TF}}(\tau)}{\mathrm{d}\tau}\right]. \tag{S37}$$

In summary, our approximant $\bar{C}_{\mathrm{TF}\gamma}(\tau)$ shall work when Eqs. (S35) and (S36) are satisfied. As we will show later, Eq. (S35) is in fact easy to satisfy and thus only Eq. (S36) tends to serve as a key factor for the validity of $\bar{C}_{\mathrm{TF}\gamma}(\tau)$.

On the other hand, the QSSA in Eq. (7) or (S21) would be valid in the following condition from Eq. (S25), instead of the condition in Eq. (S36):

$$\varepsilon_{\mathrm{TFQ}}(\tau) \equiv \frac{1}{\bar{C}_{\mathrm{TFQ}}(\tau)} \left|\frac{1}{1+\bar{A}_{\mathrm{TF}}(\tau)} \frac{\mathrm{d}\bar{C}_{\mathrm{TFQ}}(\tau)}{\mathrm{d}\tau} - \frac{1}{[1+\bar{A}_{\mathrm{TF}}(\tau)]^2} \frac{\mathrm{d}^2\bar{C}_{\mathrm{TFQ}}(\tau)}{\mathrm{d}\tau^2}\right| \ll 1, \tag{S38}$$

where $\bar{C}'_{\mathrm{TFQ}}(\tau) = \mu_{\mathrm{TF}}(\tau)\bar{C}_{\mathrm{TFQ}}(\tau)/[1 + \bar{A}_{\mathrm{TF}}(\tau)]$ from Eq. (S21) and $\bar{C}''_{\mathrm{TFQ}}(\tau)$ is given by Eq. (S37).

Regarding $\bar{C}_{\mathrm{TF}}(\tau)$ from Eqs. (S20) and (S43) in Text S8, one may expect that the high accuracy of $\bar{C}_{\mathrm{TF}\gamma}(\tau)$ in the phases might be indicated by the range of the valid conditions of $\bar{C}_{\mathrm{TF}\gamma}(\tau)$ in Eqs. (S35) and (S36). In fact, most of our simulated conditions (92.0%) in Table S3 satisfy $\max_\tau[\varepsilon_{\mathrm{TF}}(\tau)] \le 0.1$ and therefore only $\varepsilon_{\mathrm{TF}\gamma}(\tau)$ and $\varepsilon_{\mathrm{TFQ}}(\tau)$ in Eqs. (S36) and (S38) remain the key determinants of the validities of $\bar{C}_{\mathrm{TF}\gamma}(\tau)$ and $\bar{C}_{\mathrm{TFQ}}(\tau)$, respectively. Our analysis shows that $\max_\tau[\varepsilon_{\mathrm{TF}\gamma}(\tau)] \le 0.1$ for 81.6% of the conditions, slightly more than the conditions (69.9%) with $\max_\tau[\varepsilon_{\mathrm{TFQ}}(\tau)] \le 0.1$. Among them, the conditions with $\max_\tau[\varepsilon_{\mathrm{TF}\gamma}(\tau)] \le 0.1$ cover all of the conditions with $\max_\tau[\varepsilon_{\mathrm{TFQ}}(\tau)] \le 0.1$, supporting more general applicability of the ETS than the QSSA's. In addition, Fig. S1(b) shows that the ETS is particularly more valid than the QSSA for larger $K$ and smaller $k_\delta$, which tend to lengthen the effective time delay $k_\delta^{-1}[1 + K^{-1}A_{\mathrm{TF}}(t)]^{-1}$ in Eq. (8).

**Text S6. Metabolic reaction and transport kinetics**

Imagine that substrate A binds to enzyme B, which catalyzes a metabolic reaction to convert A to another molecule. The formation of the enzyme–substrate complex follows Eq. (1). In Eq. (1), we set $k_\delta = k_{\mathrm{d}} + r_{\mathrm{c}}$ where $r_{\mathrm{c}}$ is interpreted as the catalytic rate constant of the reaction (conventionally written as $k_{\mathrm{cat}}$ in literature), and $k_{\mathrm{loc}} = k_{\mathrm{dlt}} = 0$. In this system, the total substrate concentration $A(t)$ changes over time as

$$\frac{\mathrm{d}A(t)}{\mathrm{d}t} = -r_{\mathrm{c}}C(t). \quad \text{(S39)}$$

We further assume that the total enzyme concentration is constant over time [$B(t) = B$]. Eqs. (1) and (S39) fully determine the time course of the system with given initial conditions and parameters.

Unlike other molecular events that we will consider later, the majority of known metabolic reactions are likely to be modeled by the sQSSA or tQSSA to a sufficient degree, without the need for the ETS. Indeed, (*i*) most enzymatic reactions in Table S1 satisfy Eq. (S34) as well as Eqs. (S29) and (S31) (i.e., $\varepsilon_1, \varepsilon_2, \varepsilon_{\mathrm{tQ}} \ll 1$ in Table S1), although a malate dehydrogenase does not follow Eq. (S34) very well when the substrate is oxaloacetate, and (*ii*) the enzyme levels in Table S1 are generally much lower than the substrate levels, despite two exceptions of malate dehydrogenase and succinate dehydrogenase (fumarate as a substrate). These (*i*) and (*ii*) indicate that the tQSSA [relevant to (*i*)] or sQSSA [relevant to (*i*) and (*ii*)] would often suffice for the kinetic modeling of metabolic reactions. Yet, oxaloacetate conversion by malate dehydrogenase would be well described by the ETS, compared to the tQSSA and sQSSA. Indeed, the time trajectory of $C_\gamma(t)$ in Eq. (6) shows remarkable agreement with the simulated enzyme-binding substrate levels [$C(t)$] from Eqs. (1) and (S39), after some transient period of $C(t)$ that depends on the initial condition (Fig. S2); meanwhile, the tQSSA is ~0.3-ms more advanced than $C(t)$ in the overall profile and the sQSSA severely overestimates $C(t)$ [Fig. S2(b)]. For example, at $t = 1.9$ ms, $C(t) \approx C_\gamma(t) = 0.13$ $\mu$M and the sQSSA leads to 0.31 $\mu$M, while the tQSSA results in 0.13 $\mu$M at $t = 1.6$ ms [Fig. S2(b)].

The overall time shift of the tQSSA from $C(t)$ is caused by the discarding of the effective time delay in enzyme–substrate complex formation in the tQSSA. In other words, the effective time delay included in the ETS satisfactorily fixes this time shift. The overestimation of $C(t)$ by the sQSSA comes from the unlimited enzyme–substrate binding with enzyme increase $\{A(t)B(t)/[K + A(t)]\}$ in the sQSSA.

In addition, we examine the case of nutrient transport into a cell, where an "enzyme" is a transporter protein on the cell surface and a substrate is a small molecule nutrient in the extracellular environment. $r_\mathrm{c}$ is then interpreted as the uptake rate of a transporter-binding nutrient. We assume that as the cells reproduce, the transporters increase over time with constant $B'(t)/B(t)$, which is equal to the cell growth rate. With this rate, Eqs. (1) and (S39) govern the full kinetics of the nutrient transport. Our analysis of the well-documented, phosphotransferase system (PTS) in bacterium *Escherichia coli* reveals that the sQSSA alone would suffice to describe this system without the need for the ETS, as in Table S2 where $\varepsilon_1, \varepsilon_2, \varepsilon_\mathrm{tQ} \ll 1$ and the transporter level is far below the nutrient level.

**Text S7. Protein–protein interaction**

In the case of protein–protein interactions [S13–S15], the tQSSA has recently been recommended for their modeling, regarding its higher accuracy than the sQSSA's [S1,S4]. Here, we will focus on the interactions between proteins whose abundances oscillate over time with circadian rhythmicity, i.e., ~24-h periodicity. Such time-varying nature in protein abundances might challenge the relevance of the tQSSA and would serve as a testbed for the ETS.

Suppose that proteins A and B have oscillating concentrations with sinusoidal forms:

$$\overline{A}(\tau) = \overline{A}_\mathrm{max}\left\{1 - \frac{\alpha_\mathrm{A}}{2}\left[1 + \cos\left(\frac{2\pi}{k_\delta T}\tau\right)\right]\right\} \text{ and } \overline{B}(\tau) = \overline{B}_\mathrm{max}\left\{1 - \frac{\alpha_\mathrm{B}}{2}\left[1 + \cos\left(\frac{2\pi}{k_\delta T}\tau - \varphi_\mathrm{B}\right)\right]\right\}, \tag{S40}$$

where $\overline{A}(\tau)$ and $\overline{B}(\tau)$ are the dimensionless A and B concentrations in Eq. (S1), $\overline{A}_\mathrm{max}$ ($\overline{B}_\mathrm{max}$), $\alpha_\mathrm{A(B)}$, $T$, and $\varphi_\mathrm{B}$ denote the peak level of $\overline{A}(\tau)$ [$\overline{B}(\tau)$], the peak-to-trough difference of $\overline{A}(\tau)$ [$\overline{B}(\tau)$] divided by the peak level, the oscillation period of a circadian or diurnal rhythm, and the phase difference between $\overline{A}(\tau)$ and $\overline{B}(\tau)$, respectively. Here, $\alpha_\mathrm{A}$ and $\alpha_\mathrm{B}$ range from 0 to 1 (the closer they are to 1, the stronger the oscillations) and $0 \le \varphi_\mathrm{B} \le \pi$ without loss of generality. In this study, we choose $T = 24$ h. Based on $\overline{A}(\tau)$ and $\overline{B}(\tau)$ in Eq. (S40), we numerically solve Eq. (S1) to obtain $\bar{C}(\tau)$ and evaluate how well $\bar{C}(\tau)$ is approximated by the ETS in Eq. (S16), the tQSSA in Eq. (S4), and the sQSSA in Eq. (S6).

As illustrated in Fig. S3(a), we observe that the ETS tends to better match the temporal trajectory of $\bar{C}(\tau)$ than the tQSSA and sQSSA. For systematic evaluation, we define $\phi_{\gamma}^{t}$, $\phi_{\mathrm{tQ}}^{t}$, and $\phi_{\mathrm{sQ}}^{t}$ as the phase differences in hours between the ETS and $\bar{C}(\tau)$, between the tQSSA and $\bar{C}(\tau)$, and between the sQSSA and $\bar{C}(\tau)$, respectively (Text S12). The sign of a given phase difference is assigned positive (negative) if the corresponding trajectory has a more advanced (delayed) phase than $\bar{C}(\tau)$. We observe that the signs of $\phi_{\mathrm{tQ}}^{t}$ and $\phi_{\mathrm{sQ}}^{t}$ are always positive and the sign of $\phi_{\gamma}^{t}$ is mostly negative. In the example of Fig. S3(a), $\phi_{\gamma}^{t} = -0.4$ h, $\phi_{\mathrm{tQ}}^{t} = 2.4$ h, $\phi_{\mathrm{sQ}}^{t} = 2.4$ h, and hence $|\phi_{\gamma}^{t}|$ is smaller than $|\phi_{\mathrm{tQ}}^{t}|$ and $|\phi_{\mathrm{sQ}}^{t}|$. We did find that $|\phi_{\gamma}^{t}|$ tends to be smaller than $|\phi_{\mathrm{tQ}}^{t}|$ and $|\phi_{\mathrm{sQ}}^{t}|$ across physiologically-relevant conditions [Fig. S3(b), Table S3, and $P < 10^{-4}$]. Remarkably, when $|\phi_{\gamma}^{t}|$, $|\phi_{\mathrm{tQ}}^{t}|$, or $|\phi_{\mathrm{sQ}}^{t}|$ is ≥1 h, most parameter conditions (86.2%) have $|\phi_{\gamma}^{t}|$ less than both $|\phi_{\mathrm{tQ}}^{t}|$ and $|\phi_{\mathrm{sQ}}^{t}|$ at least by one hour, and some of them (22.9%) even at least by two hours [Figs. S3(c) and S3(d) and $P < 10^{-4}$].

These findings establish the tendency that the effective time delay in the ETS quantitatively well matches the phase difference between $\bar{C}(\tau)$ and its quasi-steady state. Therefore, the relaxation time in complex formation should be considered as a key to understand the deviation of the complex profile from the quasi-steady state.

Other than phases, wave profiles (determined by the waveforms and peak levels) are the important features of oscillatory molecular behaviors. Therefore, we define similarity $S_{\gamma}$ between the wave profiles of the ETS and $\bar{C}(\tau)$ by aligning their phases to the same. $S_{\gamma}$ is devised to approach 1 away from 0, as the two wave profiles quantitatively better match each other (Text S12). We also define the similarity measures $S_{\mathrm{tQ}}$ and $S_{\mathrm{sQ}}$ for the tQSSA and $\bar{C}(\tau)$, and for the sQSSA and $\bar{C}(\tau)$, respectively. In the example of Fig. S3(a), $S_{\gamma} = 0.91$, $S_{\mathrm{tQ}} = 0.90$, and $S_{\mathrm{sQ}} = 0.88$. Based on Eq. (S16), one can expect that the main difference between the ETS and tQSSA would be attributed to their phases, rather than to their shapes. As expected, $S_{\gamma}$ and $S_{\mathrm{tQ}}$ tend to be almost equal to each other [Spearman's $\rho = 0.89$ and $P < 10^{-4}$; see Fig. S3(e)]. On the other hand, consistent with the previous suggestions that the tQSSA is more accurate than the sQSSA [S1], $S_{\mathrm{sQ}}$ tends to be below $S_{\gamma}$ and $S_{\mathrm{tQ}}$ [Fig. S3(e) and $P < 10^{-4}$]. Therefore, the ETS and tQSSA better approximate the wave profile of $\bar{C}(\tau)$ than the sQSSA.

Although physiologically less relevant, the oscillatory protein levels with irregular rhythmicity may provide another testbed for the approximating capability of the ETS. Hence, we considered the following $\overline{A}(\tau)$ and $\overline{B}(\tau)$ and numerically solved Eq. (S1):

$$\overline{A}(\tau) = \frac{1}{N}\sum_{i=1}^{N}\overline{A}_{\max,i}\left\{1 - \frac{\alpha_{A,i}}{2}\left[1 + \cos\left(\frac{2\pi}{k_\delta T_{A,i}}\tau - \varphi_{A,i}\right)\right]\right\},$$

$$\overline{B}(\tau) = \frac{1}{N}\sum_{i=1}^{N}\overline{B}_{\max,i}\left\{1 - \frac{\alpha_{B,i}}{2}\left[1 + \cos\left(\frac{2\pi}{k_\delta T_{B,i}}\tau - \varphi_{B,i}\right)\right]\right\}. \quad \text{(S41)}$$

We chose $N = 10$ and randomly selected $k_\delta$ from its range in Table S3 and the other parameters from $0.1 \le \overline{A}_{\max,i}, \overline{B}_{\max,i} \le 10$, $0.5 \le \alpha_{A,i}, \alpha_{B,i} < 1$, $-\pi < \varphi_{A,i}, \varphi_{B,i} \le \pi$, and 10 h $\le T_{A,i}, T_{B,i} \le$ 40 h. We found that the ETS tends to better approximate such an irregular profile of $\overline{C}(\tau)$ than the tQSSA and sQSSA, as illustrated in Fig. S3(f).

Next, we move to a real-world example of oscillating protein interactions. In plant *Arabidopsis thaliana*, ZEITLUPE (ZTL) is an essential protein for a normal circadian periodicity. ZTL is stabilized by a direct interaction with another protein GIGANTEA (GI), and this interaction is enhanced by blue light [S16–S18]. As a result, ZTL protein levels oscillate in light–dark cycles, despite the constitutive mRNA expression of ZTL [S16]. We here assess how well the ETS accounts for the experimental ZTL profile over time, through the modeling of the ZTL–GI interaction. If $A(t)$ and $B(t)$ represent the ZTL and GI concentrations, respectively, then the ZTL turnover dynamics can be described by the following equation:

$$\frac{\mathrm{d}A(t)}{\mathrm{d}t} = g_A - r_c C(t) - r_A[A(t) - C(t)], \quad \text{(S42)}$$

where $g_A$ is the ZTL synthesis rate, $C(t)$ is the concentration of ZTL–GI complex, and $r_A$ and $r_c$ denote the degradation rates of free ZTL and GI-binding ZTL, respectively. $r_c < r_A$ because GI stabilizes ZTL. $C(t)$ is determined by Eq. (1) and we assume $k_\delta = k_d + r_c$ there. Because blue light enhances the ZTL–GI interaction, we further assume that $k_d$ and $K$ in light do not exceed $k_d$ and $K$ in darkness, respectively. We set $B(t)$ as the known GI profile [Fig. S4(a)] [S16] multiplied by a scaling coefficient $w_{GI}$, because the original GI profile is given by the concentration levels at a relative scale, not at the absolute scale. For the same reason, when comparing $A(t)$ with the experimental ZTL profile [Fig. S4(a)] [S16], we use $A_s(t) \equiv A(t)/w_{ZTL}$ where $w_{ZTL}$ is another scaling coefficient.

We compute three different versions of $A(t)$ for their comparison with the experimental ZTL profile: (*i*) the first version is the solution of $A(t)$ from Eqs. (1) and (S42), (*ii*) the second version is the solution of only Eq. (S42) with the replacement of $C(t)$ by $C_\gamma(t)$ in Eq. (6), and (*iii*) the last version is similar to version (*ii*), but with the replacement of $C(t)$ by $C_{tQ}(t)$ in Eq. (2). In other words, version (*i*) is the full modeling result that we treat as the gold standard to assess the relative accuracies of versions (*ii*) and (*iii*) from the ETS and the tQSSA, respectively. We do not consider the sQSSA because the tQSSA has already proven to be more accurate than the sQSSA in Figs. S2 and S3 as well as in previous studies [S1,S4].

We simulate the full model and its ETS and tQSSA versions in (*i*)–(*iii*) for randomly-selected parameters $g_{\mathrm{A}}$, $r_{\mathrm{A}}$, $r_{\mathrm{c}}$, $w_{\mathrm{GI}}$, and $w_{\mathrm{ZTL}}$ with $k_{\mathrm{d}}$ and $K$ in light and darkness (Table S4). We define similarity $S_{\mathrm{ZTL}}$ between $A_{\mathrm{s}}(t)$ and the empirical ZTL profile when $A_{\mathrm{s}}(t)$ is calculated from the full model by the above relation $A_{\mathrm{s}}(t) = A(t)/w_{\mathrm{ZTL}}$. $S_{\mathrm{ZTL}}$ is devised to approach 1 away from 0, as $A_{\mathrm{s}}(t)$ quantitatively better matches the ZTL profile. Analogously, $S_{\mathrm{ZTL}\gamma}$ and $S_{\mathrm{ZTLQ}}$ are defined for the ETS and tQSSA cases, respectively. Figs. S4(a) and S4(b) present an example that $A_{\mathrm{s}}(t)$ from the ETS is as close to the experimental ZTL profile as $A_{\mathrm{s}}(t)$ from the full model, and is closer to that experimental profile than $A_{\mathrm{s}}(t)$ from the tQSSA ($S_{\mathrm{ZTL}} = 0.88$, $S_{\mathrm{ZTL}\gamma} = 0.88$, and $S_{\mathrm{ZTLQ}} = 0.79$). Indeed, most of our simulated conditions (78.2%) show $S_{\mathrm{ZTL}\gamma}$ higher than $S_{\mathrm{ZTLQ}}$ [Fig. S4(c) and $P < 10^{-4}$], while $S_{\mathrm{ZTL}\gamma}$ and $S_{\mathrm{ZTL}}$ are almost the same as each other [Fig. S4(d)]. We hence conclude that the effective time delay in the ETS is important for capturing the experimental ZTL profile to a similar degree to the full modeling.

**Text S8. TF–DNA interaction**

We here consider TF–DNA interactions with circadian rhythmicity. Suppose that the TF concentration oscillates over time in a sinusoidal form:

$$\overline{A}_{\mathrm{TF}}(\tau) = \overline{A}_{\max}\left\{1 - \frac{\alpha_{\mathrm{A}}}{2}\left[1 + \cos\left(\frac{2\pi}{k_{\delta}T}\tau\right)\right]\right\}, \tag{S43}$$

where $\overline{A}_{\mathrm{TF}}(\tau)$, $\overline{A}_{\max}$, $\alpha_{\mathrm{A}}$, and $T$ are the dimensionless TF concentration in Eq. (S20), the peak level of $\overline{A}_{\mathrm{TF}}(\tau)$, the peak-to-trough difference of $\overline{A}_{\mathrm{TF}}(\tau)$ divided by the peak level, and the oscillation period of a circadian or diurnal rhythm, respectively. Here, $\alpha_{\mathrm{A}}$ ranges from 0 to 1 (the closer it is to 1, the stronger the oscillation) and we choose $T = 24$ h. Based on $\overline{A}_{\mathrm{TF}}(\tau)$ in Eq. (S43), we numerically solve Eq. (S20) to obtain $\overline{C}_{\mathrm{TF}}(\tau)$, and evaluate how well $\overline{C}_{\mathrm{TF}}(\tau)$ is approximated by the ETS in Eq. (S28) or by the QSSA in Eq. (S21).

As illustrated in Fig. S5(a), we observe that the ETS tends to better match the time trajectory of $\bar{C}_{\mathrm{TF}}(\tau)$ than the QSSA. For systematic evaluation, we define $\phi_{\gamma}^{t}$ and $\phi_{\mathrm{Q}}^{t}$ as the phase differences in hours between the ETS and $\bar{C}_{\mathrm{TF}}(\tau)$, and between the QSSA and $\bar{C}_{\mathrm{TF}}(\tau)$, respectively (Text S12). The magnitudes of these phase differences reach up to ~5 h in physiologically-relevant parameter conditions [Fig. S5(b) and Table S3]. The sign of a given phase difference is assigned positive (negative) if the corresponding trajectory has a more advanced (delayed) phase than $\bar{C}_{\mathrm{TF}}(\tau)$. In the simulated conditions (Table S3), we observe $\phi_{\mathrm{Q}}^{t} \geq 0$ and $\phi_{\gamma}^{t} \leq 0$. In the example of Fig. S5(a), $\phi_{\gamma}^{t} = -0.3$ h and $\phi_{\mathrm{Q}}^{t} = 2.3$ h, and here $|\phi_{\gamma}^{t}|$ is smaller than $|\phi_{\mathrm{Q}}^{t}|$. Indeed, our analysis suggests that $|\phi_{\gamma}^{t}|$ tends to be smaller than $|\phi_{\mathrm{Q}}^{t}|$ over the physiologically-relevant conditions [Fig. S5(b) and $P < 10^{-4}$]. When $|\phi_{\gamma}^{t}|$ or $|\phi_{\mathrm{Q}}^{t}|$ is ≥1 h, most parameter conditions (91.6%) have $|\phi_{\gamma}^{t}|$ less than $|\phi_{\mathrm{Q}}^{t}|$ at least by one hour, and a quarter of them even at least by two hours [Fig. S5(c) and $P < 10^{-4}$].

These results suggest that the effective time delay in the ETS tends to well match the phase difference between $\bar{C}_{\mathrm{TF}}(\tau)$ and its quasi-steady state. Therefore, the relaxation time in TF–DNA binding should be considered as a key to understand the deviation of $\bar{C}_{\mathrm{TF}}(\tau)$ from the quasi-steady state.

Unlike the cases of phases, we expect that the wave profiles predicted by the ETS and QSSA would be almost the same, with regards to Eq. (S28). To examine this issue, we define similarity $S_{\gamma}$ between the profiles of the ETS and $\bar{C}_{\mathrm{TF}}(\tau)$ by aligning their phases to the same. $S_{\gamma}$ is devised to approach 1 away from 0, as the two wave profiles quantitatively better match each other (Text S12). We also define the similarity $S_{\mathrm{Q}}$ for the QSSA and $\bar{C}_{\mathrm{TF}}(\tau)$. As anticipated, $S_{\gamma}$ and $S_{\mathrm{Q}}$ take almost the same values as each other (Spearman's $\rho = 0.94$ and $P < 10^{-4}$) and both are $> 0.7$ for the physiologically-relevant conditions.

Although physiologically less relevant, the oscillatory TF level with irregular rhythmicity may provide another testbed for the approximating capability of the ETS. Hence, we considered the following $\overline{A}_{\mathrm{TF}}(\tau)$ and numerically solved Eq. (S20):

$$\overline{A}_{\mathrm{TF}}(\tau) = \frac{1}{N}\sum_{i=1}^{N} \overline{A}_{\max,i} \left\{1 - \frac{\alpha_{\mathrm{A},i}}{2}\left[1 + \cos\left(\frac{2\pi}{k_{\delta} T_{\mathrm{A},i}}\tau - \varphi_{\mathrm{A},i}\right)\right]\right\}. \quad \text{(S44)}$$

We chose $N = 10$ and randomly selected $k_{\delta}$, $K$, and $V$ from the conditions in Table S3 and the other parameters from $0.1 \leq \overline{A}_{\max,i} \leq 10$, $0.5 \leq \alpha_{\mathrm{A},i} \leq 0.9$, $-\pi < \varphi_{\mathrm{A},i} \leq \pi$, and 10 h $\leq T_{\mathrm{A},i} \leq$ 40 h. Even with such irregularity of the rhythms, the ETS is still found to improve the approximation of $\bar{C}_{\mathrm{TF}}(\tau)$ compared to the QSSA, as illustrated in Fig. S5(d).

**Text S9. Positive autogenous control**

Consider a scenario of positive autoregulation that proteins enhance their own transcription after homodimer formation and this dimer–promoter interaction is facilitated by inducer molecules, as in Fig. 2(a). The protein production is hence governed by the following equations:

$$\frac{\mathrm{d}M(t)}{\mathrm{d}t} = \frac{s}{V} + (a_0 - s)C_{\mathrm{TF}}(t) - (b_0 + k_{\mathrm{dlt}})M(t), \tag{S45}$$

$$\frac{\mathrm{d}A(t)}{\mathrm{d}t} = a_1 M(t) - (r_{\mathrm{c}} + k_{\mathrm{dlt}})A(t), \tag{S46}$$

$$\frac{\mathrm{d}A_2(t)}{\mathrm{d}t} = \frac{k_{\mathrm{a}}}{2}[A(t) - 2A_2(t)]^2 - (k_{\mathrm{d}} + r_{\mathrm{c}} + k_{\mathrm{dlt}})A_2(t), \tag{S47}$$

$$\frac{\mathrm{d}C_{\mathrm{TF}}(t)}{\mathrm{d}t} = \frac{\eta\hat{k}_{\mathrm{TFa}}}{V}A_2(t) - \left[k_{\mathrm{TFd}} + k_{\mathrm{dlt}} + \eta\hat{k}_{\mathrm{TFa}}A_2(t)\right]C_{\mathrm{TF}}(t). \tag{S48}$$

Here, $M(t)$, $A(t)$, $A_2(t)$, and $C_{\mathrm{TF}}(t)$ are the total mRNA, protein, protein dimer, and promoter-binding dimer concentrations, respectively. Eqs. (S47) and (S48) are based on Eqs. (1) and (S20), but modified for homodimerization and dilution with cell growth. $s$, $a_0$, and $a_1$ denote the basal and maximal transcription rates ($s \ll a_0$) and translation rate, respectively. $b_0$, $r_{\mathrm{c}}$, and $k_{\mathrm{dlt}}$ denote the mRNA and protein degradation rates and cell growth-related dilution rate, respectively. $k_{\mathrm{a}}$, $k_{\mathrm{d}}$, and $k_{\mathrm{TFd}}$ denote the dimer association and dissociation rates and dimer–promoter dissociation rate, respectively. $\eta$ is a dimensionless quantity, monotonically increasing with an inducer level. $\eta\hat{k}_{\mathrm{TFa}}$ and $V$ correspond to $k_{\mathrm{a}}$ and $V$ in the case of Eq. (S20), respectively. We here assume that the promoter-binding dimers are neither dissociated to monomers nor degraded. To be precise, $-(r_{\mathrm{c}} + k_{\mathrm{dlt}})A(t)$ and $-(k_{\mathrm{d}} + r_{\mathrm{c}} + k_{\mathrm{dlt}})A_2(t)$ in Eqs. (S46) and (S47) should be replaced by $-r_{\mathrm{c}}[A(t) - 2C_{\mathrm{TF}}(t)] - k_{\mathrm{dlt}}A(t)$ and $-(k_{\mathrm{d}} + r_{\mathrm{c}})[A_2(t) - C_{\mathrm{TF}}(t)] - k_{\mathrm{dlt}}A_2(t)$, respectively; however, this replacement does not much alter our simulation results, and thus we keep the original forms of Eqs. (S46) and (S47) for the straightforward approximation of the dimer concentration later.

To systematically analyze the induction kinetics using dimensionless quantities, we rewrite Eqs. (S45)–(S48) as

$$\frac{\mathrm{d}\bar{M}(\hat{\tau})}{\mathrm{d}\hat{\tau}} = \sigma + (1 - \sigma)\bar{\bar{C}}_{\mathrm{TF}}(\hat{\tau}) - B_0\bar{M}(\hat{\tau}), \tag{S49}$$

$$\frac{\mathrm{d}\bar{A}(\hat{\tau})}{\mathrm{d}\hat{\tau}} = \bar{M}(\hat{\tau}) - \bar{A}(\hat{\tau}), \tag{S50}$$

$$\frac{\mathrm{d}\bar{A}_2(\hat{\tau})}{\mathrm{d}\hat{\tau}} = R[\bar{A}(\hat{\tau}) - 2\bar{A}_2(\hat{\tau})]^2 - D\bar{A}_2(\hat{\tau}), \tag{S51}$$

$$\frac{\mathrm{d}\bar{\bar{C}}_{\mathrm{TF}}(\hat{\tau})}{\mathrm{d}\hat{\tau}} = \eta P\bar{A}_2(\hat{\tau}) - [D_{\mathrm{TF}} + \eta P\bar{A}_2(\hat{\tau})]\bar{\bar{C}}_{\mathrm{TF}}(\hat{\tau}), \tag{S52}$$

where all the variables and parameters are dimensionless as $\hat{\tau} \equiv (r_{\mathrm{c}} + k_{\mathrm{dlt}})t$, $\bar{M}(\hat{\tau}) \equiv (r_{\mathrm{c}} + k_{\mathrm{dlt}})a_0^{-1}VM(t)$, $\bar{A}(\hat{\tau}) \equiv (r_{\mathrm{c}} + k_{\mathrm{dlt}})^2(a_0a_1)^{-1}VA(t)$, $\bar{A}_2(\hat{\tau}) \equiv (r_{\mathrm{c}} + k_{\mathrm{dlt}})^2(a_0a_1)^{-1}VA_2(t)$, $\bar{\bar{C}}_{\mathrm{TF}}(\hat{\tau}) \equiv VC_{\mathrm{TF}}(t)$, $\sigma \equiv sa_0^{-1} \ll 1$, $B_0 \equiv (b_0 + k_{\mathrm{dlt}})(r_{\mathrm{c}} + k_{\mathrm{dlt}})^{-1}$,

$R \equiv k_a a_0 a_1 (2V)^{-1}(r_c + k_{dlt})^{-3}$, $D \equiv (k_d + r_c + k_{dlt})(r_c + k_{dlt})^{-1}$, $P \equiv \hat{k}_{TFa} a_0 a_1 V^{-1}(r_c + k_{dlt})^{-3}$, and $D_{TF} \equiv (k_{TFd} + k_{dlt})(r_c + k_{dlt})^{-1}$.

Eqs. (S51) and (S52) are equivalent to Eqs. (S1) and (S20) with the mapping of $D\hat{\tau}$, $4RD^{-1}\bar{A}_2(\hat{\tau})$, and $2RD^{-1}\bar{A}(\hat{\tau})$ in Eq. (S51) to $\tau$, $\bar{C}(\tau)$, and both $\bar{A}(\tau)$ and $\bar{B}(\tau)$ in Eq. (S1) and that of $D_{TF}\hat{\tau}$, $\bar{\bar{C}}_{TF}(\hat{\tau})$, and $\eta P D_{TF}^{-1} \bar{A}_2(\hat{\tau})$ in Eq. (S52) to $\tau$, $KV\bar{C}_{TF}(\tau)$, and $\bar{A}_{TF}(\tau)$ in Eq. (S20), respectively. From Eqs. (S4) and (S21), the tQSSA and QSSA-based model then comprises Eq. (S50) and this equation:

$$\frac{d\bar{M}(\hat{\tau})}{d\hat{\tau}} = \sigma + (1-\sigma)\frac{\eta P D_{TF}^{-1} \bar{A}_{2tQ}(\hat{\tau})}{1+\eta P D_{TF}^{-1} \bar{A}_{2tQ}(\hat{\tau})} - B_0 \bar{M}(\hat{\tau}), \quad \text{(S53)}$$

where $\bar{A}_{2tQ}(\hat{\tau}) \equiv \left[\kappa + \bar{A}(\hat{\tau}) - \bar{\Delta}_{tQ}(\hat{\tau})\right]/2$ with $\bar{\Delta}_{tQ}(\hat{\tau}) \equiv \sqrt{\kappa[\kappa + 2\bar{A}(\hat{\tau})]}$ and $\kappa \equiv D/(4R)$. We simply call this model with Eqs. (S50) and (S53) the QSSA-based model. The second term on the right-hand side of Eq. (S53) corresponds to a TF–DNA binding curve under the quasi-steady state assumption. This binding curve is a sigmoid function of the protein level $\bar{A}(\hat{\tau})$, the known feature of positive autoregulation.

On the other hand, from Eqs. (S16) and (S28), the ETS gives rise to a model with Eq. (S50) and the following equation:

$$\frac{d\bar{M}(\hat{\tau})}{d\hat{\tau}} = \sigma + (1-\sigma)\frac{\eta P D_{TF}^{-1} \bar{A}_{2\Upsilon}\left[\hat{\tau} - \frac{1}{D_{TF} + \eta P \bar{A}_{2\Upsilon}(\hat{\tau})}\right]}{1+\eta P D_{TF}^{-1} \bar{A}_{2\Upsilon}\left[\hat{\tau} - \frac{1}{D_{TF} + \eta P \bar{A}_{2\Upsilon}(\hat{\tau})}\right]} - B_0 \bar{M}(\hat{\tau}), \quad \text{(S54)}$$

where $\bar{A}_{2\Upsilon}(\hat{\tau}) \equiv \min\left\{\bar{A}_{2tQ}\left[\hat{\tau} - R^{-1}\bar{\Delta}_{tQ}^{-1}(\hat{\tau})/4\right], \bar{A}(\hat{\tau})/2\right\}$ with the aforementioned $\bar{A}_{2tQ}(\hat{\tau})$ and $\bar{\Delta}_{tQ}(\hat{\tau})$. The second term on the right-hand side of Eq. (S54) corresponds to a TF–DNA binding curve modified with the relaxation time in the dimer–promoter interaction and dimerization.

As $\eta$ increases from 0, the simulation of the full model with Eqs. (S49)–(S52) shows that an initially low, steady-state protein level undergoes a discontinuous transition at some point $\eta = \eta_c$ [Fig. 2(b)]. Therefore, upon a sudden change of $\eta = 0$ to $\eta > \eta_c$, the protein level grows over time towards its new steady state. The induced protein time-series can be compared between the full model [Eqs. (S49)–(S52)], the ETS-based model [Eqs. (S50) and (S54)], and the QSSA-based model [Eqs. (S50) and (S53)]. These three models with the common parameter set have the same steady states, but can differ in their transient behaviors. Specifically, we will focus on the response time, defined as the time taken for a protein level to reach 90% of its steady state.

To seek the analytical expression of the response time, we consider the following condition:

$$\bar{A}(\hat{\tau}) \ll \kappa \text{ and } B_0 \gg 1, \tag{S55}$$

which allow us to take $\bar{A}_{2\mathrm{tQ}}(\hat{\tau}) \approx \bar{A}^2(\hat{\tau})/(4\kappa)$, $R^{-1}\bar{\Delta}_{\mathrm{tQ}}^{-1}(\hat{\tau})/4 \approx 1/D$, and an instantly acclimating mRNA level with $\bar{M}'(\hat{\tau}) \approx 0$ in Eqs. (S53) and (S54). Plugging them in Eq. (S50) leads to

$$\frac{\mathrm{d}\bar{A}(\hat{\tau})}{\mathrm{d}\hat{\tau}} \approx \frac{\sigma}{B_0} + \frac{1-\sigma}{B_0} \cdot \frac{\left(\frac{\eta P}{4\kappa D_{\mathrm{TF}}}\right)\bar{A}^2(\hat{\tau})}{1+\left(\frac{\eta P}{4\kappa D_{\mathrm{TF}}}\right)\bar{A}^2(\hat{\tau})} - \bar{A}(\hat{\tau}), \tag{S56}$$

$$\frac{\mathrm{d}\bar{A}(\hat{\tau})}{\mathrm{d}\hat{\tau}} \approx \frac{\sigma}{B_0} + \frac{1-\sigma}{B_0} \cdot \frac{\left(\frac{\eta P}{4\kappa D_{\mathrm{TF}}}\right)\bar{A}^2\left[\hat{\tau} - \frac{D_{\mathrm{TF}}^{-1}}{1+\left(\frac{\eta P}{4\kappa D_{\mathrm{TF}}}\right)\bar{A}^2\left(\hat{\tau}-\frac{1}{D}\right)} - \frac{1}{D}\right]}{1+\left(\frac{\eta P}{4\kappa D_{\mathrm{TF}}}\right)\bar{A}^2\left[\hat{\tau} - \frac{D_{\mathrm{TF}}^{-1}}{1+\left(\frac{\eta P}{4\kappa D_{\mathrm{TF}}}\right)\bar{A}^2\left(\hat{\tau}-\frac{1}{D}\right)} - \frac{1}{D}\right]} - \bar{A}(\hat{\tau}). \tag{S57}$$

Eqs. (S56) and (S57) come from the QSSA and ETS, respectively. $\bar{A}(\hat{\tau}) \to \sigma/B_0$ at the steady state at $\eta = 0$ and the above $\sigma \ll 1$ and $B_0 \gg 1$ ensure $\sigma/B_0 \ll 1$. Therefore, we treat the initial $\bar{A}(\hat{\tau})$ as $\sim 0$ when $\eta$ switches from 0 to $> \eta_{\mathrm{c}}$. To estimate the relevant protein response time, we consider two different stages of the protein growth: (*i*) the early stage with $0 \lesssim \bar{A}(\hat{\tau}) < 2\sqrt{\kappa D_{\mathrm{TF}}\eta^{-1}P^{-1}}$ and (*ii*) the late stage with $2\sqrt{\kappa D_{\mathrm{TF}}\eta^{-1}P^{-1}} \leq \bar{A}(\hat{\tau}) < \bar{A}_*$ where $\bar{A}_*$ is the final steady state of $\bar{A}(\hat{\tau})$. The phase portrait of Eq. (S56) in the early and late stages is illustrated by Fig. S6. For the early and late stages, Eq. (S56) respectively reduces to

$$\frac{\mathrm{d}\bar{A}(\hat{\tau})}{\mathrm{d}\hat{\tau}} \approx c + a\bar{A}^2(\hat{\tau}) - \bar{A}(\hat{\tau}) \text{ and } \frac{\mathrm{d}\bar{A}(\hat{\tau})}{\mathrm{d}\hat{\tau}} \approx \frac{1}{B_0} - \bar{A}(\hat{\tau}), \tag{S58}$$

where $a \equiv \eta P(1-\sigma)/(4\kappa B_0 D_{\mathrm{TF}})$ and $c \equiv \sigma/B_0$. Eq. (S58) gives the dimensionless form of the QSSA-based response time, as follows:

$$\begin{aligned}[\hat{\tau}]_0^{\bar{A}(\hat{\tau})=\zeta\bar{A}_*} &= \int_0^{2\sqrt{\frac{\kappa D_{\mathrm{TF}}}{\eta P}}} \frac{\mathrm{d}\bar{A}(\hat{\tau})}{\bar{A}'(\hat{\tau})} + \int_{2\sqrt{\frac{\kappa D_{\mathrm{TF}}}{\eta P}}}^{\zeta\bar{A}_*} \frac{\mathrm{d}\bar{A}(\hat{\tau})}{\bar{A}'(\hat{\tau})} \\ &\approx \frac{2}{\sqrt{\frac{\eta-\eta_*}{\eta_*}}}\left\{\arctan\left[\frac{\sqrt{\frac{(1-\sigma)\eta}{\sigma\eta_*}}-1}{\sqrt{\frac{\eta-\eta_*}{\eta_*}}}\right] + \arctan\left(\frac{1}{\sqrt{\frac{\eta-\eta_*}{\eta_*}}}\right)\right\} + \ln\left[\frac{1-2\sqrt{\frac{(1-\sigma)\sigma\eta_*}{\eta}}}{1-\zeta}\right],\end{aligned} \tag{S59}$$

where $\bar{A}_* \approx 1/B_0$ from Eq. (S58), $\zeta = 0.9$ by our definition of the response time, and

$$\eta_* \equiv \frac{B_0^2\kappa D_{\mathrm{TF}}}{P\sigma(1-\sigma)}. \tag{S60}$$

When $\eta \to \eta_*$, the response time in Eq. (S59) diverges. $\eta_*$ approximates the transition point, i.e., $\eta_\mathrm{c} \approx \eta_*$. Applying this fact and $\sigma \ll 1$ to Eq. (S59) for $\eta \to \eta_\mathrm{c}$ informs the QSSA-based response time near the transition point, as follows:

$$\frac{2\pi}{\sqrt{\frac{\eta-\eta_\mathrm{c}}{\eta_\mathrm{c}}}} + \ln\left(\frac{1-2\sqrt{\sigma}}{1-\zeta}\right). \tag{S61}$$

Note that the first diverging term roots in the early stage of the protein growth and the second term in the late stage. The above estimate tends to match the QSSA model simulation with Eqs. (S50) and (S53) under the condition of Eq. (S55) [9.6 ± 9.0% relative error (avg. ± s.d. in simulated conditions) and $P < 10^{-4}$].

Next, regarding the ETS, Eq. (S57) in the early stage of the protein growth reduces to

$$\frac{\mathrm{d}\bar{A}(\hat{\tau})}{\mathrm{d}\hat{\tau}} \approx c + a\bar{A}^2\left(\hat{\tau} - \frac{1}{D} - \frac{1}{D_\mathrm{TF}}\right) - \bar{A}(\hat{\tau}), \tag{S62}$$

where $a$ and $c$ are the same as Eq. (S58). Roughly, $\bar{A}\left(\hat{\tau} - D^{-1} - D_\mathrm{TF}^{-1}\right) > 0$ only if $\bar{A}(\hat{\tau}) > \bar{A}_\mathrm{min} \equiv c\left[1 - \left(1 + D^{-1} + D_\mathrm{TF}^{-1}\right)^{-1}\right]$, because $\bar{A}\left(\hat{\tau} - D^{-1} - D_\mathrm{TF}^{-1}\right) \approx \bar{A}(\hat{\tau}) - \left(D^{-1} + D_\mathrm{TF}^{-1}\right)\bar{A}'(\hat{\tau})$ and $\bar{A}'(\hat{\tau})$ is given by

$$\frac{\mathrm{d}\bar{A}(\hat{\tau})}{\mathrm{d}\hat{\tau}} \approx \frac{2a\left(D^{-1}+D_\mathrm{TF}^{-1}\right)\bar{A}(\hat{\tau})+1-\sqrt{4a\left(D^{-1}+D_\mathrm{TF}^{-1}\right)^2\left[\left(1+\frac{1}{D^{-1}+D_\mathrm{TF}^{-1}}\right)\bar{A}(\hat{\tau})-c\right]+1}}{2a\left(D^{-1}+D_\mathrm{TF}^{-1}\right)^2}. \tag{S63}$$

$\bar{A}'(\hat{\tau})$ above is the solution of $a\left(D^{-1} + D_\mathrm{TF}^{-1}\right)^2[\bar{A}'(\hat{\tau})]^2 - \left[2a\left(D^{-1} + D_\mathrm{TF}^{-1}\right)\bar{A}(\hat{\tau}) + 1\right]\bar{A}'(\hat{\tau}) + a\bar{A}^2(\hat{\tau}) - \bar{A}(\hat{\tau}) + c \approx 0$. It comes from Eq. (S62) and $\bar{A}^2\left(\hat{\tau} - D^{-1} - D_\mathrm{TF}^{-1}\right) \approx \bar{A}^2(\hat{\tau}) - 2\left(D^{-1} + D_\mathrm{TF}^{-1}\right)\bar{A}(\hat{\tau})\bar{A}'(\hat{\tau}) + \left(D^{-1} + D_\mathrm{TF}^{-1}\right)^2[\bar{A}'(\hat{\tau})]^2$ as $\bar{A}''(\hat{\tau})\sim 0$ in the early growth stage when $\eta$ is not too larger than $\eta_\mathrm{c}$ so that $\bar{A}(\hat{\tau})$ exhibits prolonged slow growth. Apart from Eq. (S63) for $\bar{A}(\hat{\tau}) > \bar{A}_\mathrm{min}$ in the early growth stage, we obtain $\bar{A}'(\hat{\tau}) \approx c - \bar{A}(\hat{\tau})$ for $\bar{A}(\hat{\tau}) \leq \bar{A}_\mathrm{min}$ by setting $\bar{A}\left(\hat{\tau} - D^{-1} - D_\mathrm{TF}^{-1}\right)$ in Eq. (S62) as $\sim 0$. $\bar{A}'(\hat{\tau})$ in the late stage of the protein growth is the same as the second equation in Eq. (S58), i.e., not different from the QSSA. Therefore, the response time from the ETS takes this form:

$$[\hat{\tau}]_0^{\bar{A}(\hat{\tau})=\zeta\bar{A}_*} = \int_0^{\bar{A}_{\min}} \frac{\mathrm{d}\bar{A}(\hat{\tau})}{\bar{A}'(\hat{\tau})} + \int_{\bar{A}_{\min}}^{2\sqrt{\frac{\kappa D_{\mathrm{TF}}}{\eta P}}} \frac{\mathrm{d}\bar{A}(\hat{\tau})}{\bar{A}'(\hat{\tau})} + \int_{2\sqrt{\frac{\kappa D_{\mathrm{TF}}}{\eta P}}}^{\zeta\bar{A}_*} \frac{\mathrm{d}\bar{A}(\hat{\tau})}{\bar{A}'(\hat{\tau})}$$

$$\approx \ln\left(1+\frac{1}{D}+\frac{1}{D_{\mathrm{TF}}}\right) + \left(\frac{1}{D}+\frac{1}{D_{\mathrm{TF}}}\right)\ln\left\{1+\frac{DD_{\mathrm{TF}}(u-1)[DD_{\mathrm{TF}}(u-1)-2(D+D_{\mathrm{TF}})]}{(D+D_{\mathrm{TF}})^2\frac{\eta}{\eta_*}}\right\}$$

$$+\frac{2}{\sqrt{\frac{\eta-\eta_*}{\eta_*}}}\left(1+\frac{1}{D}+\frac{1}{D_{\mathrm{TF}}}\right)\left\{\arctan\left[\frac{DD_{\mathrm{TF}}(u-1)-D-D_{\mathrm{TF}}}{(D+D_{\mathrm{TF}})\sqrt{\frac{\eta-\eta_*}{\eta_*}}}\right]+\arctan\left(\frac{1}{\sqrt{\frac{\eta-\eta_*}{\eta_*}}}\right)\right\}+\ln\left[\frac{1-2\sqrt{\frac{(1-\sigma)\sigma\eta_*}{\eta}}}{1-\zeta}\right],$$

(S64)

where $u \equiv \sqrt{\sqrt{\eta\eta_*^{-1}}(D^{-1}+D_{\mathrm{TF}}^{-1})^2\left\{2[1+(D^{-1}+D_{\mathrm{TF}}^{-1})^{-1}]\sqrt{\sigma^{-1}(1-\sigma)}-\sqrt{\eta\eta_*^{-1}}\right\}+1}$ and the other notations are the same as Eq. (S59).

In a similar fashion to the previous trial, we put $\eta_{\mathrm{c}} \approx \eta_*$ and $\sigma \ll 1$ into Eq. (S64) when $\eta \to \eta_{\mathrm{c}}$. The ETS then predicts the near-transition response time as follows:

$$\frac{2\pi}{\sqrt{\frac{\eta-\eta_{\mathrm{c}}}{\eta_{\mathrm{c}}}}}\left(1+\frac{1}{D}+\frac{1}{D_{\mathrm{TF}}}\right)+\ln\left(\frac{1-2\sqrt{\sigma}}{1-\zeta}\right)+\ln\left(1+\frac{1}{D}+\frac{1}{D_{\mathrm{TF}}}\right)$$
$$+\left(\frac{1}{D}+\frac{1}{D_{\mathrm{TF}}}\right)\ln\left\{1+\frac{DD_{\mathrm{TF}}(\bar{u}-1)[DD_{\mathrm{TF}}(\bar{u}-1)-2(D+D_{\mathrm{TF}})]}{(D+D_{\mathrm{TF}})^2}\right\}, \quad \text{(S65)}$$

where $\bar{u} \equiv \sqrt{(D^{-1}+D_{\mathrm{TF}}^{-1})^2\{2[1+(D^{-1}+D_{\mathrm{TF}}^{-1})^{-1}]\sigma^{-1/2}-1\}+1}$. Remarkably, the full model simulation with Eqs. (S49)–(S52) under the condition of Eq. (S55) does support the above predicted response time [14.3 ± 12.6% relative error (avg. ± s.d. in simulated conditions) and $P < 10^{-4}$]. Although the predicted response time looks complex in its form, only the first term $2\pi\left(1+D^{-1}+D_{\mathrm{TF}}^{-1}\right)/\sqrt{(\eta-\eta_{\mathrm{c}})/\eta_{\mathrm{c}}}$ becomes dominating as $\eta \to \eta_{\mathrm{c}}$. $D^{-1}+D_{\mathrm{TF}}^{-1}$ in the term represents the unique feature of the ETS, as will be discussed below.

Subtracting Eq. (S61) from Eq. (S65) suggests that the exact response time would be longer than the QSSA-based estimate by

$$\frac{2\pi}{\sqrt{\frac{\eta-\eta_{\mathrm{c}}}{\eta_{\mathrm{c}}}}}\left(\frac{1}{D}+\frac{1}{D_{\mathrm{TF}}}\right)+\ln\left(1+\frac{1}{D}+\frac{1}{D_{\mathrm{TF}}}\right)+\left(\frac{1}{D}+\frac{1}{D_{\mathrm{TF}}}\right)\ln\left\{1+\frac{DD_{\mathrm{TF}}(\bar{u}-1)[DD_{\mathrm{TF}}(\bar{u}-1)-2(D+D_{\mathrm{TF}})]}{(D+D_{\mathrm{TF}})^2}\right\}. \quad \text{(S66)}$$

The dimensional form of Eq. (S66) is given by Eq. (9) with a notation $r \equiv r_{\mathrm{c}} + k_{\mathrm{dlt}}$. This result was verified by the full and QSSA model simulation difference, as discussed in the main text.

The above difference between the exact and QSSA-based response times originates in the early stage of the protein growth, as evident from a comparison between Eqs. (S59) and (S64). This response time difference vanishes as $D^{-1}+D_{\mathrm{TF}}^{-1} \to 0$. Because $D^{-1}$ and $D_{\mathrm{TF}}^{-1}$ are proportional to the effective time delays in dimerization and dimer–promoter interaction, respectively, the total effective time delay $D^{-1} + D_{\mathrm{TF}}^{-1}$ is responsible for the retarded response compared to the QSSA. Strikingly, the response time difference diverges as $\eta \to \eta_{\mathrm{c}}$. This relatively far retarded response near the transition point is an amplified effect of the effective time delay (i.e., the relaxation time in complex formation), attributed to the ultrasensitivity of the protein growth around the trough of the near-transition phase portrait in Fig. S6.

**Text S10. Negative autogenous control**

Consider a scenario of negative autoregulation that proteins repress their own transcription after homodimer formation and this dimer–promoter interaction is inhibited by inducer molecules, as in Fig. S7(a). In fact, the homodimerization is not essential for our main results later, but just considered for a fair comparison of this system and the above positive autoregulation case. The protein production process is described by the following equations and Eqs. (S46) and (S47):

$$\frac{\mathrm{d}M(t)}{\mathrm{d}t} = \frac{s}{V} + (a_0 - s)\left[\frac{1}{V} - C_{\mathrm{TF}}(t)\right] - (b_0 + k_{\mathrm{dlt}})M(t), \tag{S67}$$

$$\frac{\mathrm{d}C_{\mathrm{TF}}(t)}{\mathrm{d}t} = \frac{\hat{k}_{\mathrm{TFa}}}{\eta V} A_2(t) - \left[k_{\mathrm{TFd}} + k_{\mathrm{dlt}} + \frac{\hat{k}_{\mathrm{TFa}}}{\eta} A_2(t)\right] C_{\mathrm{TF}}(t), \tag{S68}$$

where all the variables and parameters are the same as Eqs. (S45) and (S48) and $\hat{k}_{\mathrm{TFa}}/\eta$ corresponds to $k_{\mathrm{a}}$ in the case of Eq. (S20). As the simulated inducer level increases with $\eta$, Fig. S7(b) demonstrates that the steady-state protein level $[A(t)]$ increases, but does not show a discrete transition like the positive regulation case. Upon acute induction by an $\eta = 0$ to $\eta > 0$ switch, the protein level grows to the new steady state over time and this response becomes slower at larger $\eta$ [Fig. S7(c)]. Still, the response is speedier than in the positive regulation case, when the protein steady states are similar in that comparison [Figs. 2(c) and S7(c)]—consistent with the previous finding of the beneficial effect of negative autoregulation [S19].

Adopting the dimensionless variables and parameters in Eqs. (S49) and (S52), we rewrite Eqs. (S67) and (S68) as

$$\frac{\mathrm{d}\bar{M}(\hat{\tau})}{\mathrm{d}\hat{\tau}} = \sigma + (1 - \sigma)\left[1 - \bar{\bar{C}}_{\mathrm{TF}}(\hat{\tau})\right] - B_0 \bar{M}(\hat{\tau}), \tag{S69}$$

$$\frac{\mathrm{d}\bar{\bar{C}}_{\mathrm{TF}}(\hat{\tau})}{\mathrm{d}\hat{\tau}} = \frac{P}{\eta}\bar{A}_2(\hat{\tau}) - \left[D_{\mathrm{TF}} + \frac{P}{\eta}\bar{A}_2(\hat{\tau})\right]\bar{\bar{C}}_{\mathrm{TF}}(\hat{\tau}). \tag{S70}$$

Eq. (S70) is equivalent to Eq. (S20) with the mapping of $D_{\mathrm{TF}}\hat{\tau}$, $\bar{\bar{C}}_{\mathrm{TF}}(\hat{\tau})$, and $P\eta^{-1}D_{\mathrm{TF}}^{-1}\bar{A}_2(\hat{\tau})$ to $\tau$, $KV\bar{C}_{\mathrm{TF}}(\tau)$, and $\bar{A}_{\mathrm{TF}}(\tau)$, respectively. Therefore, based on the same procedure as the positive regulation case, we obtain the QSSA-based model with Eq. (S50) and the following equation:

$$\frac{\mathrm{d}\bar{M}(\hat{\tau})}{\mathrm{d}\hat{\tau}} = \sigma + \frac{(1-\sigma)\eta}{\eta + PD_{\mathrm{TF}}^{-1}\bar{A}_{2\mathrm{tQ}}(\hat{\tau})} - B_0\bar{M}(\hat{\tau}). \tag{S71}$$

Likewise, the ETS leads to the model with Eq. (S50) and the following equation:

$$\frac{\mathrm{d}\bar{M}(\hat{\tau})}{\mathrm{d}\hat{\tau}} = \sigma + \frac{(1-\sigma)\eta}{\eta + PD_{\mathrm{TF}}^{-1}\bar{A}_{2\gamma}\left[\hat{\tau} - \frac{\eta}{\eta D_{\mathrm{TF}} + P\bar{A}_{2\gamma}(\hat{\tau})}\right]} - B_0\bar{M}(\hat{\tau}). \tag{S72}$$

$\bar{A}_{2\mathrm{tQ}}(\hat{\tau})$ and $\bar{A}_{2\gamma}(\hat{\tau})$ in Eqs. (S71) and (S72) are defined as in Eqs. (S53) and (S54).

When there is a sudden $\eta = 0$ to $\eta > 0$ change, the response time of the induced protein can be compared between the full model [Eqs. (S50), (S51), (S69), and (S70)], the ETS model [Eqs. (S50) and (S72)], and the QSSA model [Eqs. (S50) and (S71)]. Across physiologically-relevant conditions in Table S5, we found that the ETS model tends to better agree with the full model in the response time than the QSSA model, especially for small $\eta$ values as exemplified by Fig. S7(c) [$P < 10^{-4}$; in Fig. S7(c) (left), the ETS even reproduces an overshoot in the protein level, whereas the QSSA does not]. Like the positive regulation case, we will try the analytical formulation of the response times with the ETS and the QSSA.

We assume Eq. (S55) for Eqs. (S50), (S71), and (S72) and obtain the following expressions with a notation $\eta_{\mathrm{s}} \equiv P/(4\kappa D_{\mathrm{TF}}B_0^2)$:

$$\frac{\mathrm{d}\bar{A}(\hat{\tau})}{\mathrm{d}\hat{\tau}} \approx \frac{\sigma}{B_0} + \frac{1-\sigma}{B_0}\cdot\frac{1}{1+\frac{\eta_{\mathrm{s}}}{\eta}B_0^2\bar{A}^2(\hat{\tau})} - \bar{A}(\hat{\tau}), \tag{S73}$$

$$\frac{\mathrm{d}\bar{A}(\hat{\tau})}{\mathrm{d}\hat{\tau}} \approx \frac{\sigma}{B_0} + \frac{1-\sigma}{B_0}\cdot\frac{1}{1+\frac{\eta_{\mathrm{s}}}{\eta}B_0^2\bar{A}^2\left[\hat{\tau} - \frac{D_{\mathrm{TF}}^{-1}}{1+\frac{\eta_{\mathrm{s}}}{\eta}B_0^2\bar{A}^2\left(\hat{\tau}-\frac{1}{D}\right)} - \frac{1}{D}\right]} - \bar{A}(\hat{\tau}). \tag{S74}$$

Eqs. (S73) and (S74) pertain to the QSSA and ETS, respectively. The phase portrait of Eq. (S73) in Fig. S7(d) suggests that the closer to the steady state, the more rate-limiting in the protein response process. We therefore focus on the late-stage dynamics and assume

$$\bar{A}(\hat{\tau}) \approx \bar{A}_* - \epsilon e^{-\lambda(\hat{\tau}-\hat{\tau}_0)}, \tag{S75}$$

where $\bar{A}_*$ is the steady state of $\bar{A}(\hat{\tau})$, $\lambda$ and $\epsilon$ are positive constants, and $\hat{\tau}_0$ denotes the initial point of $\hat{\tau}$. Eq. (S75) would work for large $\eta$ without an overshoot in $\bar{A}(\hat{\tau})$. We first set $\bar{A}^2(\hat{\tau} - D^{-1})$ in Eq. (S74) as $\sim\bar{A}_*^2$ and then apply Eq. (S75) to Eq. (S74). Consequently,

$$\epsilon\lambda e^{-\lambda(\hat{\tau}-\hat{\tau}_0)} \approx \frac{\sigma}{B_0} + \frac{1-\sigma}{B_0} \cdot \frac{1}{1+\frac{\eta_\mathrm{S}}{\eta} B_0^2 \left[\bar{A}_* - \epsilon e^{-\lambda(\hat{\tau}-\hat{\tau}_\mathrm{d}-\hat{\tau}_0)}\right]^2} + \epsilon e^{-\lambda(\hat{\tau}-\hat{\tau}_0)} - \bar{A}_*, \qquad \text{(S76)}$$

where $\hat{\tau}_\mathrm{d} \equiv D_\mathrm{TF}^{-1}[1 + \eta^{-1}\eta_\mathrm{S} B_0^2 \bar{A}_*^2]^{-1} + D^{-1}$. The Taylor expansion of Eq. (S76) up to $O(\epsilon)$ gives rise to

$$\frac{2\eta_\mathrm{S} B_0 \bar{A}_* (1-\sigma)}{\eta\left(1+\frac{\eta_\mathrm{S}}{\eta} B_0^2 \bar{A}_*^2\right)^2} e^{\lambda\hat{\tau}_\mathrm{d}} - \lambda + 1 \approx 0. \qquad \text{(S77)}$$

The use of relation $e^x \geq 1 + x$ for Eq. (S77) with $x = \lambda\hat{\tau}_\mathrm{d}$ leads to

$$\lambda \gtrsim \frac{\eta\left(1+\frac{\eta_\mathrm{S}}{\eta} B_0^2 \bar{A}_*^2\right)^2 + 2\eta_\mathrm{S} B_0 \bar{A}_* (1-\sigma)}{\eta\left(1+\frac{\eta_\mathrm{S}}{\eta} B_0^2 \bar{A}_*^2\right)^2 - 2\hat{\tau}_\mathrm{d}\eta_\mathrm{S} B_0 \bar{A}_* (1-\sigma)}. \qquad \text{(S78)}$$

In a similar manner to the positive regulation case, we treat the initial $\bar{A}(\hat{\tau})$ as $\sim 0$. Compatibly, we treat $\bar{A}(\hat{\tau} - \hat{\tau}_\mathrm{d})$ as $\sim 0$ for $\hat{\tau} \leq \hat{\tau}_0 + \hat{\tau}_\mathrm{d}$, as well. In that time period, $\bar{A}'(\hat{\tau}) \approx B_0^{-1} - \bar{A}(\hat{\tau})$ from the modified Eq. (S74), and therefore $\bar{A}(\hat{\tau}_0 + \hat{\tau}_\mathrm{d}) \approx B_0^{-1}\left(1 - e^{-\hat{\tau}_\mathrm{d}}\right)$. For the continuity of this result with Eq. (S75) at $\hat{\tau} \to \hat{\tau}_0 + \hat{\tau}_\mathrm{d}$, it should be satisfied that $\epsilon \approx B_0^{-1} e^{(\lambda-1)\hat{\tau}_\mathrm{d}} - (B_0^{-1} - \bar{A}_*) e^{\lambda\hat{\tau}_\mathrm{d}}$. With this form of $\epsilon$ and the quantity $\zeta$ used for Eqs. (S59) and (S64), we solve Eq. (S75) for $\hat{\tau} - \hat{\tau}_0$ when $\bar{A}(\hat{\tau}) = \zeta\bar{A}_*$, and this value of $\hat{\tau} - \hat{\tau}_0$ is the estimated response time in a dimensionless form. For simplicity, if one focuses on the case of $\eta \gg \eta_\mathrm{S}$, $\bar{A}_* \approx B_0^{-1}$ from Eq. (S73) or (S74). Combining the current procedure with the relation in Eq. (S78) suggests the upper limit of the ETS-based response time, as follows:

$$\ln\left(\frac{1}{1-\zeta}\right) - 2(1-\sigma)\left(1 + \frac{1}{D} + \frac{1}{D_\mathrm{TF}}\right)\left[\ln\left(\frac{1}{1-\zeta}\right) - \frac{1}{D} - \frac{1}{D_\mathrm{TF}}\right]\frac{\eta_\mathrm{S}}{\eta}, \qquad \text{(S79)}$$

which is obtained by the Taylor expansion up to $O(\eta_\mathrm{S}/\eta)$ for $\eta \gg \eta_\mathrm{S}$. Although Eq. (S79) intends to be the upper limit, it is in practice close to the simulated response time from the full model with Eqs. (S50), (S51), (S69), and (S70) under the condition of Eq. (S55) [2.8 ± 1.6% relative error (avg. ± s.d. in simulated conditions) and $P < 10^{-4}$].

Following the above way, one can also calculate the QSSA-based response time with Eq. (S73). In Eq. (S78), one should replace $\hat{\tau}_\mathrm{d}$ by zero and use ≈ instead of ≳. When $\eta \gg \eta_\mathrm{S}$, the QSSA-based response time up to $O(\eta_\mathrm{S}/\eta)$ is given by

$$\ln\left(\frac{1}{1-\zeta}\right)\left[1 - 2(1-\sigma)\frac{\eta_\mathrm{S}}{\eta}\right]. \qquad \text{(S80)}$$

This estimate tends to match the QSSA model simulation with Eqs. (S50) and (S71) under the condition of Eq. (S55) [2.8 ± 1.7% relative error (avg. ± s.d. in simulated conditions) and $P < 10^{-4}$].

Subtracting Eq. (S79) from Eq. (S80) informs the lower limit of the response time difference between the QSSA-based estimate and our prediction, as follows:

$$2(1-\sigma)\left(\frac{1}{D}+\frac{1}{D_{\mathrm{TF}}}\right)\left[\ln\left(\frac{1}{1-\zeta}\right)-1-\frac{1}{D}-\frac{1}{D_{\mathrm{TF}}}\right]\frac{\eta_{\mathrm{s}}}{\eta}. \quad \text{(S81)}$$

This minimum difference approaches zero when $D^{-1}+D_{\mathrm{TF}}^{-1}\to 0$. Because $D^{-1}$ and $D_{\mathrm{TF}}^{-1}$ are proportional to the effective time delays in dimerization and dimer–promoter interaction, respectively, the total effective time delay $D^{-1}+D_{\mathrm{TF}}^{-1}$ is responsible for the advanced response with the retarded inhibition of transcription.

Dividing Eq. (S81) by $r_{\mathrm{c}}+k_{\mathrm{dlt}}$ gives the dimensional form of Eq. (S81). Consistent with our analytical prediction, the QSSA-to-full model difference in their simulated response times is linearly scaled to $\eta_{\mathrm{s}}/\eta$ ($R^2>0.87$) and its slope against $\eta_{\mathrm{s}}/\eta$ equals or exceeds that in the dimensional form of Eq. (S81) for most of the simulated conditions [88.8%; see Fig. S7(e)]. In the example of Fig. S7(e), the QSSA model does overestimate the response time by about ten minutes and the error diminishes for larger $\eta$ values.

**Text S11. Rhythmic protein degradation**

To understand the rhythmicity in the experimental degradation rates of circadian proteins, we constructed the kinetic model of circadian protein production and degradation. Here, $A_0(t)$ and $A_1(t)$ represent the concentrations of unmodified and modified proteins, respectively, while the protein modified by ubiquitination undergoes degradation. The protein turnover dynamics is described by

$$\frac{\mathrm{d}A_0(t)}{\mathrm{d}t}=g(t)-a_0A_0(t), \quad \text{(S82)}$$

$$\frac{\mathrm{d}A_1(t)}{\mathrm{d}t}=a_0A_0(t)-r_{\mathrm{c}}A_1(t), \quad \text{(S83)}$$

where $g(t)$ and $a_0$ are the protein synthesis (translation) and modification rates, respectively, and $r_{\mathrm{c}}$ is the modified protein's turnover rate. If the protein turnover requires multiple preceding PTMs like mono- or multisite phosphorylation and subsequent ubiquitination, we consider Eq. (S82) and the following equation instead of Eq. (S83):

$$\frac{\mathrm{d}A_i(t)}{\mathrm{d}t}=a_{i-1}A_{i-1}(t)-a_iA_i(t), \quad \text{(S84)}$$

where $A_i(t)$ denotes the concentration of the $i$-th modified protein with $i=1,2,\cdots,n$ ($n$ is the total number of the PTMs), $a_i$ for $i\le n-1$ denotes the rate of the $(i+1)$-th modification, and $a_n\equiv r_{\mathrm{c}}$, the turnover rate of the $n$-th modified protein. In the case of $n=1$, Eq.

(S84) becomes the same as Eq. (S83). Therefore, we will consider Eqs. (S82) and (S84) whether $n = 1$ or $n > 1$ [these equations are identical to Eqs. (10) and (11) in the main text].

For the total protein concentration $A(t) \equiv \sum_{i=0}^{n} A_i(t)$, Eqs. (S82) and (S84) result in

$$\frac{\mathrm{d}A(t)}{\mathrm{d}t} = g(t) - r(t)A(t), \tag{S85}$$

where $r(t)$ is the protein degradation rate given by

$$r(t) = r_\mathrm{c}\frac{A_n(t)}{A(t)}. \tag{S86}$$

To exclude the possibility of the time-dependent regulation of the degradation process, $a_i$'s with $0 \le i \le n$ in Eqs. (S82) and (S84) are constants. Given the circadian profile of protein synthesis rate $g(t)$, the numerical solution of Eqs. (S82), (S84), and (S86) gives rise to the degradation rate $r(t)$. In this calculation, we use the sinusoidal form of $g(t)$:

$$g(t) = g_\mathrm{max}\left\{1 - \frac{\alpha_\mathrm{g}}{2}\left[1 + \cos\left(\frac{2\pi t}{T}\right)\right]\right\}, \tag{S87}$$

where $g_\mathrm{max}$, $\alpha_\mathrm{g}$, and $T$ are constants.

On the other hand, the ETS provides the analytical estimate of $r(t)$ through the comparison of seemingly unrelated but mathematically equivalent systems. Because the periodic $g(t)$ assures $\langle A_i'(t)\rangle = \langle A_i(t)\rangle' = 0$ ($\langle\cdot\rangle$ is a time average), $\langle g(t)\rangle = a_i\langle A_i(t)\rangle$ and $\langle A_i(t)\rangle$ is then inversely proportional to $a_i$. Hence, $A(t) \approx A_u(t) + A_v(t)$ for $i = u, v$ ($u < v$) that hold the two smallest $a_i$ values among all $a_i$'s. In addition, $a_i$'s are larger for $i \neq u, v$ by definition and therefore the corresponding $A_i(t)$'s are likely to follow $A_i'(t) \approx 0$ with the effect of $-a_iA_i(t)$ in Eq. (S84). Subsequently, $a_vA_v(t) \approx a_{v+1}A_{v+1}(t) \approx \cdots \approx r_\mathrm{c}A_n(t)$. As a result, Eqs. (S84) and (S86) reduce to

$$\begin{aligned}\frac{\mathrm{d}A_v(t)}{\mathrm{d}t} &\approx \sum_{i=u+1}^{v}\frac{\mathrm{d}A_i(t)}{\mathrm{d}t} = a_uA_u(t) - a_vA_v(t)\\ &\approx a_u[A(t) - A_v(t)] - a_vA_v(t),\end{aligned} \tag{S88}$$

$$r(t) \approx a_v\frac{A_v(t)}{A(t)}. \tag{S89}$$

When $n = 1$, it is obvious that $u = 0$, $v = 1$, and symbol = replaces ≈ in Eqs. (S88) and (S89). Eqs. (S88) and (S89) may not satisfactorily work for large $n$ due to accumulating errors in the approximation, but still capture the core structure of the dynamics.

We then observe the mathematical equivalence of Eqs. (S20) and (S88) despite their different biological contexts: $C_\mathrm{TF}(t)$ and $A_v(t)$, $V^{-1}$ and $A(t)$, $k_\mathrm{a}A_\mathrm{TF}(t)$ and $a_u$, and $k_\delta$ and $a_v$ in correspondence. Based on this correspondence and Eqs. (7), (8), and (S89) with the condition $A_v(t) \le A(t)$, the ETS gives the estimate of $r(t)$ as $r_\gamma(t)$ in Eq. (12). Following a similar procedure, the QSSA gives the estimate of $r(t)$ as $a_u a_v/(a_u + a_v)$. The ETS-estimated degradation rate $r_\gamma(t)$ is rhythmic when the protein level $A(t)$ is rhythmic, but the QSSA-based degradation rate is just constant over time.

**Text S12. Simulation and analysis methods**

***Overview***

Numerical simulations and analyses were performed by Python 3.7.0 or 3.7.4. Ordinary differential equations (ODEs) were solved by LSODA (scipy.integrate.solve_ivp) in SciPy v1.1.0 or v1.3.1 with the maximum time step of 0.05 h. Delay differential equations were solved by a modified version of the ddeint module with LSODA [S20].

Splines of discrete data points were achieved with scipy.interpolate.splrep in SciPy v1.3.1. Linear regression of data points was performed with scipy.stats.linregress in SciPy v1.3.1 and then the slope of the fitted line and $R^2$ were obtained.

For the parameter selection in numerical simulations or for the null model generation in statistical significance tests, random numbers were sampled by the Mersenne Twister in random.py.

Spearman's $\rho$ was measured with scipy.stats.spearmanr in SciPy v1.1.0 or v1.3.1. To test the significance of $\rho$ between two groups of variables, we randomized the pairing of the variables between the groups (while maintaining the original group membership) and measured the $P$ value (one-tailed) from the $10^4$ null configurations. This method was also applied to the significance test of the average of the relative errors of analytical estimates against actual simulation data in Texts S9 and S10.

***Malate dehydrogenase system in Text S6***

The numerical solution of $A(t_1)$ from Eqs. (1) and (S39) $\{t_1 \equiv t - [k_\delta \Delta_{\mathrm{tQ}}(t)]^{-1}\}$ is needed to obtain $C_\gamma(t)$ in Eq. (6) through the calculation of $C_{\mathrm{tQ}}(t_1)$. However, that solution at $t_1$ may not be available for given time $t$ because of a finite single time step in ODE solving. Therefore, we took the solution at time $t_2$ which is earlier than but the closest to $t_1$ among the available time points, and set this solution as the initial condition to solve again Eqs. (1) and (S39) from time $t_2$ to $t_1$. We used the resulting solution $A(t_1)$ to calculate $C_{\mathrm{tQ}}(t_1)$ and then $C_\gamma(t)$.

***Protein–protein and TF–DNA interaction models in Texts S7 and S8***

$10^5$ parameter sets were randomly selected for Eqs. (S1) and (S40) or for Eqs. (S20) and (S43), according to Table S3. We found that the simulated $\bar{C}(\tau)$ and $\bar{C}_{\mathrm{TF}}(\tau)$ were insensitive to their initial conditions. A phase difference between two periodic time-series was calculated by maximizing their cross-correlation with a varying displacement of one series relative to the other [S21]. For the cross-correlation calculation, the time average of each series was shifted to zero and ten duplicates of a single time period ($10 \times T$) was used. The cross-correlation was obtained with signal.correlate in SciPy v1.1.0 or v1.3.1 (mode = 'same' and method = 'fft'). In the case of irregular oscillation simulations, $10^2$ parameter sets were randomly selected for Eqs. (S1) and (S41) [Eqs. (S20) and (S44)] from the ranges noted below Eq. (S41) [Eq. (S44)]. To test the significance of the observation in Text S7 that $|\phi_\gamma^t|$ tends to be smaller than $|\phi_{\mathrm{tQ}}^t|$, we obtained $\Lambda = \mathrm{med}(|\phi_{\mathrm{tQ}}^t|) - \mathrm{med}(|\phi_\gamma^t|)$ [where $\mathrm{med}(\cdot)$ is the median over parameter sets] and randomized the labeling of $\bar{C}_\gamma(\tau)$ and the tQSSA [$\bar{C}_{\mathrm{tQ}}(\tau)$] for each parameter set. We then measured $\Lambda_{\mathrm{r}} = \mathrm{med}(|\phi_{\mathrm{tQ}}^t|) - \mathrm{med}(|\phi_\gamma^t|)$ with the new sets of $|\phi_{\mathrm{tQ}}^t|$ and $|\phi_\gamma^t|$ in this null configuration. The $P$ value was given by the probability of $\Lambda_{\mathrm{r}} \geq \Lambda$ across $10^4$ null configurations. The analogous methods were applied to the cases of $|\phi_\gamma^t|$ vs. $|\phi_{\mathrm{sQ}}^t|$, $S_\gamma$ vs. $S_{\mathrm{sQ}}$, and $S_{\mathrm{tQ}}$ vs. $S_{\mathrm{sQ}}$ in Text S7, and $|\phi_\gamma^t|$ vs. $|\phi_{\mathrm{Q}}^t|$ in Text S8. In Text S7, we tested the significance of the fraction ($q$) of parameter sets with $|\phi_\gamma^t| < \min(\phi_{\mathrm{tQ}}^t, \phi_{\mathrm{sQ}}^t) - \upsilon$ ($\upsilon = 1$h or 2h) as follows, when $|\phi_\gamma^t|$, $|\phi_{\mathrm{tQ}}^t|$, or $|\phi_{\mathrm{sQ}}^t|$ is ≥1h: for each parameter set, we randomized the labeling of $\bar{C}_\gamma(\tau)$, the tQSSA, and the sQSSA, and measured the fraction ($q_{\mathrm{r}}$) of the parameter sets of $|\phi_\gamma^t| < \min(\phi_{\mathrm{tQ}}^t, \phi_{\mathrm{sQ}}^t) - \upsilon$ with the new $|\phi_\gamma^t|$, $|\phi_{\mathrm{tQ}}^t|$, and $|\phi_{\mathrm{sQ}}^t|$ when $|\phi_\gamma^t|$, $|\phi_{\mathrm{tQ}}^t|$, or $|\phi_{\mathrm{sQ}}^t|$ is ≥1h. The $P$ value was given by the probability of $q_{\mathrm{r}} \geq q$ across $10^4$ null configurations. The analogous method was applied to the case of $|\phi_\gamma^t|$ vs. $|\phi_{\mathrm{Q}}^t|$ in Text S8.

We measured the similarity between two time-series $f_1(t)$ and $f_2(t)$ as $\left\{\int_t^{t+T} \min[f_1(t'), f_2(t')]dt'\right\} / \int_t^{t+T} \max[f_1(t'), f_2(t')]dt'$, where $T$ is an oscillation period of $f_1(t)$ and $f_2(t)$ [S22]. This quantity takes a range of 0 to 1, and becomes large for quantitatively similar profiles of $f_1(t)$ and $f_2(t)$. In the cases of $S_\gamma$, $S_{\mathrm{tQ}}$, and $S_{\mathrm{sQ}}$ in Text S7 and $S_\gamma$ and $S_{\mathrm{Q}}$ in Text S8, the phase difference between two time-series in the comparison was set to zero by the phase shift of the one series to the other, before measuring their profile similarity.

***ZTL–GI interaction in Text S7***

We achieved the cubic splines of the experimental GI and ZTL levels in equal length light–dark cycles [S23] (data originally from Ref. [S16]). For ZTL profile simulation, we multiplied this GI spline by a scaling coefficient $w_{\mathrm{GI}}$, and used this profile as $B(t)$. From Table S4, $10^5$ parameter sets were randomly selected for the full model and its ETS and tQSSA versions. We found that the simulated ZTL profile from the full model was insensitive to the initial condition of $C(t)$. To test the significance of the fraction ($q$) of parameter sets with $S_{\mathrm{ZTL}\gamma} > S_{\mathrm{ZTLQ}}$, we randomized the labeling of the ETS and tQSSA results for each parameter set and measured the fraction ($q_{\mathrm{r}}$) of the parameter sets of $S_{\mathrm{ZTL}\gamma} > S_{\mathrm{ZTLQ}}$ with the new $S_{\mathrm{ZTL}\gamma}$ and $S_{\mathrm{ZTLQ}}$. The $P$ value was given by the probability of $q_{\mathrm{r}} \geq q$ across $10^4$ null configurations.

***Positive and negative autoregulation in Texts S9 and S10***

**(associated with Section *Autogenous control* in the main text)**

$10^5$ and 500 parameter sets were randomly selected for simulation without and with the condition of Eq. (S55), respectively, as described in Table S5. The simulation results were insensitive to the initial conditions of variables. The steady state of a protein level was heuristically identified by the conditions of $|\bar{A}(\hat{\tau}_1) - \bar{A}(\hat{\tau}_1 - 1)| < |\bar{A}(\hat{\tau}_1 - 1) - \bar{A}(\hat{\tau}_1 - 2)|$ and $\max[\bar{A}(\hat{\tau}_1), \bar{A}(\hat{\tau}_1 - 1), \bar{A}(\hat{\tau}_1 - 2)] - \min[\bar{A}(\hat{\tau}_1), \bar{A}(\hat{\tau}_1 - 1), \bar{A}(\hat{\tau}_1 - 2)] < 0.01 \min[\bar{A}(\hat{\tau}_1), \bar{A}(\hat{\tau}_1 - 1), \bar{A}(\hat{\tau}_1 - 2)]$: we considered $\bar{A}(\hat{\tau}_1)$ in these conditions as the steady state of $\bar{A}(\hat{\tau})$. The steady states of other variables were determined in a similar way, too. To ensure $A_2(t), A(t) - 2A_2(t) \gg V^{-1}$ (at least at the steady state) for the validity of Eqs. (S47) and (S48) [or (S68)], we first obtained the analytical solution of the steady states of $A(t)$ and $A_2(t)$ from a model without autoregulation [$M'(t) = sV^{-1} + \eta(\eta+1)^{-1}(a_0 - s)V^{-1} - (b_0 + k_{\mathrm{dlt}})M(t)$ and Eqs. (S46) and (S47)] and used these rough estimates to select only the parameters with $A_2(t), A(t) - 2A_2(t) > 10V^{-1}$; after this crude initial screening, the actual simulation results with the correct model were considered only when they satisfied $A_2(t), A(t) - 2A_2(t) > 10V^{-1}$ at the steady states.

The protein response time was defined as the first time of the protein level to cross 90% of the steady state with the initial condition of $\bar{A}(\hat{\tau}_0) = \sigma/B_0$ ($\sigma/B_0$ is the exact steady state for $\eta = 0$ at $\hat{\tau} < \hat{\tau}_0$). For the simulated response times from the full model ($Z$), the ETS-based model ($X$), and the QSSA-based model ($Y$), we tested the significance of the fraction ($q$) of parameter sets with $|(X-Z)/Z| < |(Y-Z)/Z|$ as follows, when $|(X-Z)/Z|$ or $|(Y-Z)/Z|$ is >0.1: for each parameter set, we randomized the labeling of $X$ and $Y$, and measured the fraction ($q_{\mathrm{r}}$) of the parameter sets of $|(X-Z)/Z| < |(Y-Z)/Z|$ with the new $X$ and $Y$ when $|(X-Z)/Z|$ or $|(Y-Z)/Z|$ is >0.1. The $P$ value was given by the probability of $q_{\mathrm{r}} \geq q$ across $10^4$ null configurations. On the other hand, in the case of positive autoregulation, the full model simulation showed that the transition point $\eta = \eta_{\mathrm{c}}$ was very close to $\eta_*$ in Eq. (S60) within a range of $1.05\eta_* \leq \eta_{\mathrm{c}} \leq 1.1\eta_*$. Therefore, we identified $\eta_{\mathrm{c}}$ by the full model simulation across 1,000 grid points of $\eta$ from $1.05\eta_*$ to $1.1\eta_*$—the right point of the abrupt increase of the response time, more than twice the immediate one. To compare the linear regression of simulation data to Eq. (S66) [equivalently, Eq. (9) in its dimensional form], $\eta$ was uniformly sampled from the following range for each parameter set in the condition of Eq. (S55): $1.1\eta_c \leq \eta \leq 2\eta_c$. We selected this range to consider $\eta$ close to $\eta_c$, but not too close because of a small difference between $\eta_*$ and $\eta_\mathrm{c}$ as $1.05\eta_* \leq \eta_\mathrm{c} \leq 1.1\eta_*$. To compare the linear regression of simulation data to Eq. (S81), $\eta$ was uniformly sampled from the following range for each parameter set in the condition of Eq. (S55): $10\eta_\mathrm{s} \leq \eta \leq 100\eta_\mathrm{s}$. We selected this range to consider $\eta \gg \eta_\mathrm{s}$.

***Rhythmic protein degradation in Text S11***

**(associated with Section *Rhythmic degradation of circadian proteins* in the main text)**

$10^5$ parameter sets were randomly selected for Eqs. (S82), (S84), (S86), and (S87) as described in Table S6. To make a fair comparison of simulation results between different $n$'s, we selected the same $g_\mathrm{max}$, $B$, $r_\mathrm{c}$, and $k_{n-i}$ (for given $i$, $i = 1, 2, \cdots$) across different $n$'s. We found that the simulated $A(t)$ was insensitive to the initial conditions. Although $A(t_1)$ [$t_1 \equiv t - (a_u + a_v)^{-1}$] is needed for the calculation of $r_\gamma(t)$ in Eq. (12), it may not be available for a given moment $t$ because of a finite single time step in ODE solving. Therefore, we used the cubic spline of the solved $A(t)$ to estimate $A(t_1)$, and this estimated $A(t_1)$ was almost identical to the higher time-resolution solution of the ODEs at time $t_1$. This interpolation method saves the computational time compared to the ODE solving with higher time resolution. Regarding Table S6, we considered $A(t)$ as periodic if the minimum-to-maximum difference of the peak levels over three periods [$3 \times T$ with $T$ in Eq. (S87)] was less than 1% of the minimum peak level. The peak times of $r(t)$ and $-A'(t)/A(t)$ were averaged over three periods. To compare the probability distributions of $r(t)$'s relative amplitude (or its estimate) from different $n$'s, we used the simulation results with similar $A(t)$ profiles: for a given value of $x$, we chose the simulation results with $A(t)$'s relative amplitude between $x - 0.05$ and $x + 0.05$ [e.g., $x = 1$ in the case of Fig. 3(e)].

***Section Parameter estimation in the main text***

We considered protein–protein interactions with time-varying protein concentrations $A(t)$ and $B(t)$ from Eq. (S40) by $A(t) = K\overline{A}(\tau)$, $B(t) = K\overline{B}(\tau)$, and $t = k_\delta^{-1}\tau$. Likewise, we considered TF–DNA interactions with time-varying TF concentration $A_\mathrm{TF}(t)$ from Eq. (S43) by $A_\mathrm{TF}(t) = K\overline{A}_\mathrm{TF}(\tau)$ and $t = k_\delta^{-1}\tau$. We randomly sampled 25,000 parameter sets from Table S3. Given the $A(t)$ and $B(t)$ profiles [$A_\mathrm{TF}(t)$ profile] and sampled parameter set, $C(t)$ [$C_\mathrm{TF}(t)$] was determined by Eq. (1) [Eq. (S20)]. A series of $C(t)$ or $C_\mathrm{TF}(t)$ at 408h $\leq t \leq$ 480h every two hours [$C(0) = C_{\mathrm{TF}}(0) = 0$] was used as a "true" dataset for the estimation of parameters $K$ and $k_\delta$. We then fitted the ETS [Eq. (6)], tQSSA [Eq. (2)], or sQSSA [Eq. (5)] to those series of $C(t)$ by minimizing the mean squared error. Similarly, we fitted the ETS [Eq. (8)] or QSSA [Eq. (7)] to $C_{\mathrm{TF}}(t)$. For the error minimization, we applied Powell's method [S24] (scipy.optimize.minimize) in SciPy v1.5.2. Alternatively, we also tried Trust Region Reflective algorithm [S25] (scipy.optimize.least_squares) in SciPy v1.5.2 for the fitting of the tQSSA and QSSA, but this method did not much change the estimated parameters. During the error minimization, the ranges of $K$ and $k_\delta$ were constrained as 0.01 nM $\leq K \leq$ 1,000 nM and 0.1 $\mathrm{h}^{-1} \leq k_\delta \leq$ 10 $\mathrm{h}^{-1}$, consistent with Table S3. For each parameter estimation, we tried ten initial conditions of the parameters and considered the output parameters with the smallest error among these ten cases.

### References

S1. J. K. Kim and J. J. Tyson, *Misuse of the Michaelis–Menten Rate Law for Protein Interaction Networks and Its Remedy*, PLoS Comput. Biol. **16**, e1008258 (2020).

S2. J. A. M. Borghans, R. J. de Boer, and L. A. Segel, *Extending the Quasi-Steady State Approximation by Changing Variables*, Bull. Math. Biol. **58**, 43–63 (1996).

S3. A. R. Tzafriri, *Michaelis–Menten Kinetics at High Enzyme Concentrations*, Bull. Math. Biol. **65**, 1111–1129 (2003).

S4. J. K. Kim and D. B. Forger, *A Mechanism for Robust Circadian Timekeeping via Stoichiometric Balance*, Mol. Syst. Biol. **8**, 630 (2012).

S5. L. A. Segel and M. Slemrod, *The Quasi-Steady-State Assumption: a Case Study in Perturbation*, SIAM Rev. **31**, 446–477 (1989).

S6. V. Henri, *Lois Générales de l'Action des Diastases* (Librairie Scientifique A. Hermann, 1903).

S7. L. Michaelis and M. L. Menten, *Die Kinetik der Invertinwirkung*, Biochem. Z. **49**, 333–369 (1913).

S8. G. E. Briggs and J. B. S. Haldane, *A Note on the Kinetics of Enzyme Action*, Biochem. J. **19**, 338 (1925).

S9. J. Gunawardena, *Time-Scale Separation–Michaelis and Menten's Old Idea, Still Bearing Fruit*, FEBS. J. **281**, 473–488 (2014).

S10. V. Kampen and N. Godfried, *Stochastic Processes in Physics and Chemistry* (Elsevier, 1992).

S11. J. K. Kim and E. D. Sontag, *Reduction of Multiscale Stochastic Biochemical Reaction Networks Using Exact Moment Derivation*, PLOS Comput. Biol. **13**, e1005571 (2017).

S12. Y. M. Song, H. Hong, and J. K. Kim, *Universally Valid Reduction of Multiscale Stochastic Biochemical Systems Using Simple Non-Elementary Propensities*, PLOS Comput. Biol. **17**, e1008952 (2021).

S13. A. Fujioka, K. Terai, R. E. Itoh, K. Aoki, T. Nakamura, S. Kuroda *et al.*, *Dynamics of the Ras/ERK MAPK Cascade as Monitored by Fluorescent Probes*, J. Biol. Chem. **281**, 8917–8926 (2006).

S14. N. Blüthgen, F. J. Bruggeman, S. Legewie, H. Herzel, H. V. Westerhoff, and B. N. Kholodenko, *Effects of Sequestration on Signal Transduction Cascades*, FEBS J. **273**, 895–906 (2006).

S15. S. Carmi, E. Y. Levanon, S. Havlin, and E. Eisenberg, *Connectivity and Expression in Protein Networks: Proteins in a Complex Are Uniformly Expressed*, Phys. Rev. E. **73**, 031909 (2006).

S16. W.-Y. Kim, S. Fujiwara, S.-S. Suh, J. Kim, Y. Kim, L. Han *et al*., *ZEITLUPE is a Circadian Photoreceptor Stabilized by GIGANTEA in Blue Light*, Nature **449**, 356–360 (2007).

S17. J. Y. Cha, J. Kim, T.-S. Kim, Q. Zeng, L. Wang, S. Y. Lee, W.-Y. Kim, and D. E. Somers, *GIGANTEA Is a Co-Chaperone Which Facilitates Maturation of ZEITLUPE in the Arabidopsis Circadian Clock*, Nat. Commun. **8**, 3 (2017).

S18. C.-M. Lee, M.-W. Li, A. Feke, W. Liu, A. M. Saffer, and J. M. Gendron, *GIGANTEA Recruits the UBP12 and UBP13 Deubiquitylases to Regulate Accumulation of the ZTL Photoreceptor Complex*, Nat. Commun. **10**, 3750 (2019).

S19. N. Rosenfeld, M. B. Elowitz, and U. Alon, *Negative Autoregulation Speeds the Response Times of Transcription Networks*, J. Mol. Biol. **323**, 785–793 (2002).

S20. Scipy-based delay differential equation (dde) solver, https://github.com/Zulko/ddeint.

S21. H. Wang and P. Chu, V*oice Source Localization for Automatic Camera Pointing System in Video Conferencing*, 1997 IEEE International Conference on Acoustics, Speech, and Signal Processing **1** , 187–190 (1997).

S22. R. Lim, J. Chae, D. E. Somers, C.-M. Ghim, and P.-J. Kim, *Cost-Effective Circadian Mechanism: Rhythmic Degradation of Circadian Proteins Spontaneously Emerges without Rhythmic Post-Translational Regulation*, iScience **24**, 102726 (2021).

S23. M. Foo, D. E. Somers, and P.-J. Kim, *Kernel Architecture of the Genetic Circuitry of the Arabidopsis Circadian System*, PLoS Comput. Biol. **12**, e1004748 (2016).

S24. M. J. D. Powell, *An Efficient Method for Finding the Minimum of a Function of Several Variables without Calculating Derivatives*, Comput. J. **7**, 155–162 (1964).

S25. M. A. Branch, T. F. Coleman, and Y. Li, *A Subspace, Interior, and Conjugate Gradient Method for Large-Scale Bound-Constrained Minimization Problems*, SIAM J. Sci. Comput. **21**, 1 (1999).

S26. K. R. Albe, M. H. Butler, and B. E. Wright, *Cellular Concentrations of Enzymes and Their Substrates*, J. Theor. Biol. **143**, 163 (1990).

S27. B. D. Bennett, E. H. Kimball, M. Gao, R. Osterhout, S. J. V. Dien, and J. D. Rabinowitz, *Absolute Metabolite Concentrations and Implied Enzyme Active Site Occupancy in Escherichia coli*, Nat. Chem. Biol. **5**, 593 (2009).

S28. A. Chang, L. Jeske, S. Ulbrich, J. Hofmann, J. Koblitz, I. Schomburg, M. Neumann-Schaal, D. Jahn, and D. Schomburg, *BRENDA, the ELIXIR Core Data Resource in 2021: New Developments and Updates*, Nucleic Acids Res. **49**, D498 (2021).

S29. J. O. Park, S. A. Rubin, Y.-F. Xu, D. Amador-Noguez, J. Fan, T. Shlomi, and J. D. Rabinowitz, *Metabolite Concentrations, Fluxes and Free Energies Imply Efficient Enzyme Usage*, Nat. Chem. Biol. **12**, 482 (2016).

S30. J. M. Rohwer, N. D. Meadow, S. Roseman, H. V. Westerhoff, and P. W. Postma, *Understanding Glucose Transport by the Bacterial Phosphoenolpyruvate: Glycose Phosphotransferase System on the Basis of Kinetic Measurements in Vitro*, J. Biol. Chem. **275**, 34909 (2000).

S31. K. B. Andersen and K. von Meyenburg, *Are Growth Rates of Escherichia coli in Batch Cultures Limited by Respiration?*, J. Bacteriol. **144**, 114 (1980).

S32. J. M. Berg, J. L. Tymoczko, G. J. Gatto Jr., and L. Stryer, *Biochemistry* (W. H. Freeman, 2015).

S33. A. Bassal *et al.*, *Reshaping of the Arabidopsis thaliana Proteome Landscape and Co-Regulation of Proteins in Development and Immunity*, Mol. Plant **13**, 1709 (2020).

S34. J. Chae, R. Lim, C.-M. Ghim, and P.-J. Kim, *Backward Simulation for Inferring Hidden Biomolecular Kinetic Profiles*, STAR Protoc. **2**, 100958 (2021).

S35. H.-H. Jo, Y. J. Kim, J. K. Kim, M. Foo, D. E. Somers, and P.-J. Kim, *Waveforms of Molecular Oscillations Reveal Circadian Timekeeping Mechanisms*, Commun. Biol. **1**, 207 (2018).

S36. S. Proshkin, A. R. Rahmouni, A. Mironov, and E. Nudler, *Cooperation between Translating Ribosomes and RNA Polymerase in Transcription Elongation*, Science **328**, 504 (2010).

S37. J. A. Bernstein, A. B. Khodursky, P.-H. Lin, S. Lin-Chao, and S. N. Cohen, *Global Analysis of mRNA Decay and Abundance in Escherichia coli at Single-Gene Resolution Using Two-Color Fluorescent DNA Microarrays*, Proc. Natl. Acad. Sci. USA **99**, 9697 (2002).

S38. A. Arkin, J. Ross, and H. H. McAdams, *Stochastic Kinetic Analysis of Developmental Pathway Bifurcation in Phage λ-Infected Escherichia coli Cells*, Genetics **149**, 1633 (1998).

S39. H. Chen, K. Shiroguchi, H. Ge, and X. S. Xie, *Genome-Wide Study of mRNA Degradation and Transcript Elongation in Escherichia coli*, Mol. Syst. Biol. **11**, 781 (2015).

S40. M. R. Maurizi, *Proteases and Protein Degradation in Escherichia coli*, Experientia **48**, 178 (1992).

S41. K. Robison, A. M. McGuire, and G. M. Church, *A Comprehensive Library of DNA-Binding Site Matrices for 55 Proteins Applied to the Complete Escherichia coli K-12 Genome*, J. Mol. Biol. **284**, 241 (1998).

S42. H. E. Kubitschek and J. A. Friske, *Determination of Bacterial Cell Volume with the Coulter Counter*, J. Bacteriol. **168**, 1466 (1986).

S43. R. Narumi, Y. Shimizu, M. Ukai-Tadenuma, K. L. Ode, G. N. Kanda, Y. Shinohara, A. Sato, K. Matsumoto, and H. R. Ueda, *Mass Spectrometry-Based Absolute Quantification Reveals Rhythmic Variation of Mouse Circadian Clock Proteins*, Proc. Natl. Acad. Sci. USA **113**, E3461 (2016).

S44. M. Y. Hein *et al.*, *A Human Interactome in Three Quantitative Dimensions Organized by Stoichiometries and Abundances*, Cell **163**, 712 (2015).

### Figures and Tables

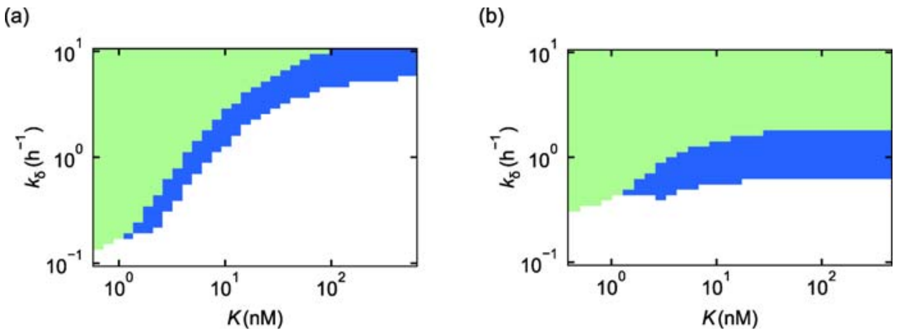


**Fig. S1. Preconditions of rate laws.** (a) The ranges of $K$ and $k_\delta$ valid for the ETS with $\max_\tau[\varepsilon_1(\tau)] \leq 0.1$, $\max_\tau[\varepsilon_2(\tau)] \leq 0.1$, and $\max_\tau\left[\varepsilon_\gamma(\tau)\right] \leq 0.1$ cover the ranges for the tQSSA with $\max_\tau\left[\varepsilon_{\text{tQ}}(\tau)\right] \leq 0.1$ instead of $\max_\tau\left[\varepsilon_\gamma(\tau)\right] \leq 0.1$. Green represents the ranges common for both the ETS and tQSSA, and blue represents those only for the ETS. The calculations are based on Eqs. (S4), (S5), (S29), (S31), (S32), (S34), and (S40) (Texts S5 and S7). (b) In the case of TF–DNA interactions, the ranges of $K$ and $k_\delta$ valid for the ETS with $\max_\tau[\varepsilon_{\text{TF}}(\tau)] \leq 0.1$ and $\max_\tau\left[\varepsilon_{\text{TF}\gamma}(\tau)\right] \leq 0.1$ cover the ranges for the QSSA with $\max_\tau\left[\varepsilon_{\text{TFQ}}(\tau)\right] \leq 0.1$ instead of $\max_\tau\left[\varepsilon_{\text{TF}\gamma}(\tau)\right] \leq 0.1$. Green represents the ranges common for both the ETS and QSSA, and blue represents those only for the ETS. The calculations are based on Eqs. (S21), (S35), (S36), (S38), and (S43) (Texts S5 and S8). In (a) and (b), parameters are selected from Table S3, and their specific values are presented in Table S10.

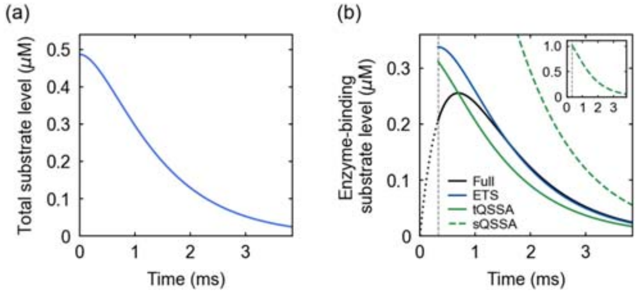


**Fig. S2. Oxaloacetate (substrate) conversion by malate dehydrogenase (enzyme).** (a) The total substrate concentration over time, calculated by the full model of Eqs. (1) and (S39). (b) The enzyme-binding substrate concentrations from the full model (black dotted line for a transient period described below), ETS, tQSSA, and sQSSA. These calculations are based on the total substrate concentration in (a). An inset shows a more complete range of the enzyme-binding substrate concentration from the sQSSA. In (a) and (b), we used the parameters and total enzyme concentration in Table S1 and set the initial substrate concentration as the substrate concentration in Table S1. When solving Eqs. (1) and (S39), the initial concentration of the enzyme-binding substrate was set to zero. As discussed in Text S1, any rate law without the initial-condition dependency would work only for $t \gg \left[k_\delta \Delta_{\mathrm{tQ}}(0)\right]^{-1}$, and also the ETS is ill-defined for a period $t < \left[k_\delta \Delta_{\mathrm{tQ}}(t)\right]^{-1}$; therefore, for the right comparison with the ETS, (b) presents the tQSSA and sQSSA results only after $t = \left[k_\delta \Delta_{\mathrm{tQ}}(t)\right]^{-1}$ (vertical dashed line).

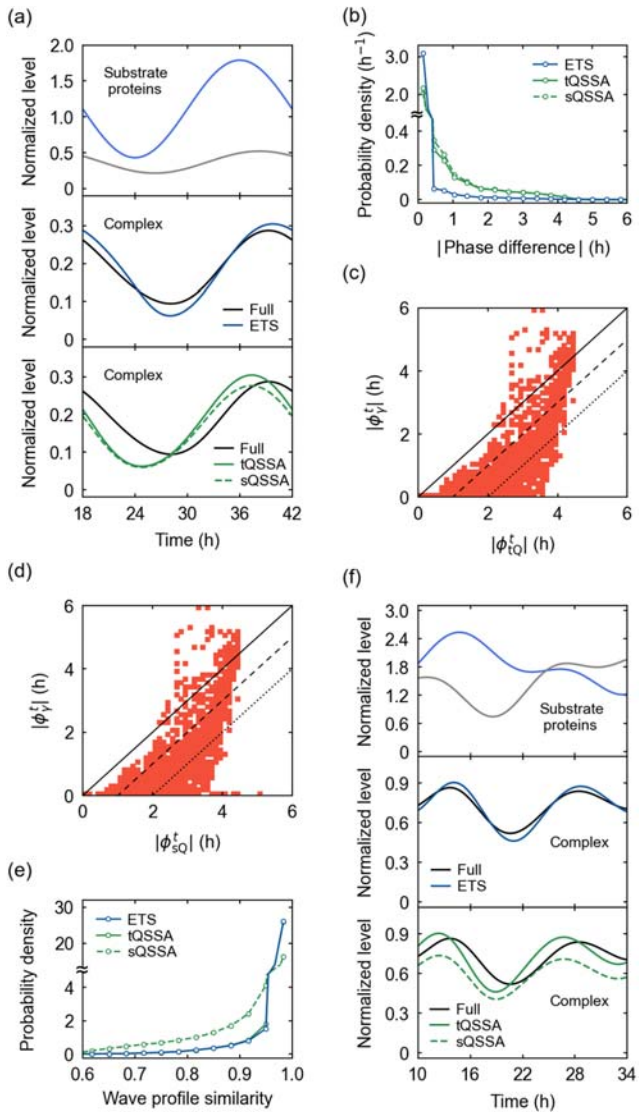


**Fig. S3. Protein–protein interaction modeling.** (a) Example time-series of substrate protein levels $\bar{A}(\tau)$ and $\bar{B}(\tau)$ in Eq. (S40) at the top, the full model-based complex level $\bar{C}(\tau)$ and the ETS at the center, and $\bar{C}(\tau)$, the tQSSA, and the sQSSA at the bottom. $t = k_\delta^{-1}\tau$ as defined before Eq. (S1). (b) Probability distributions of $|\phi_\gamma^t|$ ("ETS"), $|\phi_{\mathrm{tQ}}^t|$ ("tQSSA"), and $|\phi_{\mathrm{sQ}}^t|$ ("sQSSA") over randomly-sampled parameter sets in Table S3. (c,d) Scatter plot of $|\phi_{\mathrm{tQ}}^t|$ and $|\phi_\gamma^t|$ (c), or that of $|\phi_{\mathrm{sQ}}^t|$ and $|\phi_\gamma^t|$ (d), when $|\phi_\gamma^t|$, $|\phi_{\mathrm{tQ}}^t|$, or $|\phi_{\mathrm{sQ}}^t|$ is ≥1 h with randomly-sampled parameter sets in Table S3. A solid diagonal line corresponds to $|\phi_\gamma^t| = |\phi_{\mathrm{tQ}}^t|$ (c) or $|\phi_\gamma^t| = |\phi_{\mathrm{sQ}}^t|$ (d), a dashed diagonal line to $|\phi_\gamma^t| = |\phi_{\mathrm{tQ}}^t| - 1\mathrm{h}$ (c) or $|\phi_\gamma^t| = |\phi_{\mathrm{sQ}}^t| - 1\mathrm{h}$ (d), and a dotted diagonal line to $|\phi_\gamma^t| = |\phi_{\mathrm{tQ}}^t| - 2\mathrm{h}$ (c) or $|\phi_\gamma^t| = |\phi_{\mathrm{sQ}}^t| - 2\mathrm{h}$ (d). Although not covered in (b) and (d), $|\phi_{\mathrm{sQ}}^t| > 6\mathrm{h}$ for a tiny portion of the parameter sets (0.03%), in which still $|\phi_\gamma^t|, |\phi_{\mathrm{tQ}}^t| \leq 1\mathrm{h}$. (e) Probability distributions of $S_\gamma$ ("ETS"), $S_{\mathrm{tQ}}$ ("tQSSA"), and $S_{\mathrm{sQ}}$

(“sQSSA”) over randomly-sampled parameter sets in Table S3. (f) Example time-series of substrate protein levels $\bar{A}(\tau)$ and $\bar{B}(\tau)$ with irregular rhythmicity in Eq. (S41) at the top, the full model-based complex level $\bar{C}(\tau)$ and the ETS at the center, and $\bar{C}(\tau)$, the tQSSA, and the sQSSA at the bottom. For more details of (a)–(f), refer to Text S7 and Tables S11 and S12.

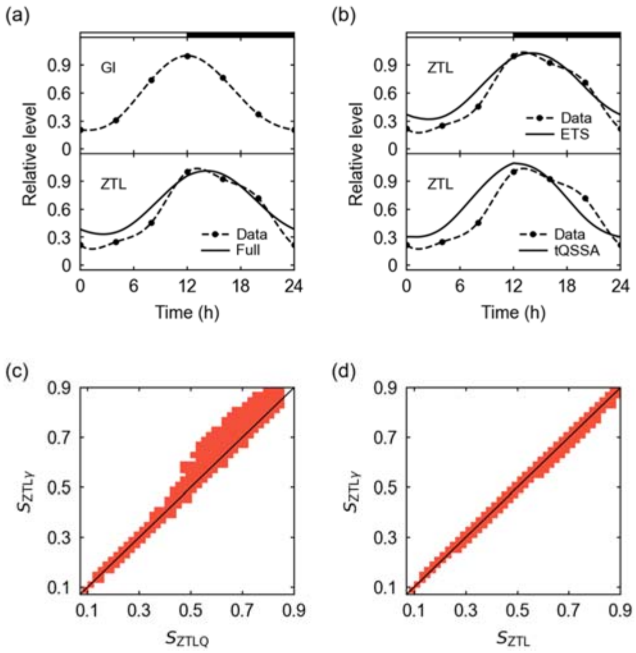


**Fig. S4. Protein ZTL–GI interaction in *Arabidopsis*.** (a) The experimental GI levels [S16] and their interpolation at the top, and the experimental ZTL levels [S16], their interpolation, and the full model-simulated ZTL profile at the bottom. (b) The experimental ZTL profile in (a), together with the ETS-based profile at the top and the tQSSA-based profile at the bottom. In (a) and (b), horizontal white and black segments correspond to light and dark intervals, respectively. For model parameters in (a) and (b), refer to Table S13. (c,d) Scatter plot of $S_{\mathrm{ZTLQ}}$ and $S_{\mathrm{ZTL}\gamma}$ (c), or that of $S_{\mathrm{ZTL}}$ and $S_{\mathrm{ZTL}\gamma}$ (d), over randomly-selected parameter sets in Table S4. A diagonal line corresponds to $S_{\mathrm{ZTL}\gamma} = S_{\mathrm{ZTLQ}}$ (c) or $S_{\mathrm{ZTL}\gamma} = S_{\mathrm{ZTL}}$ (d). For more details of (a)–(d), refer to Text S7.

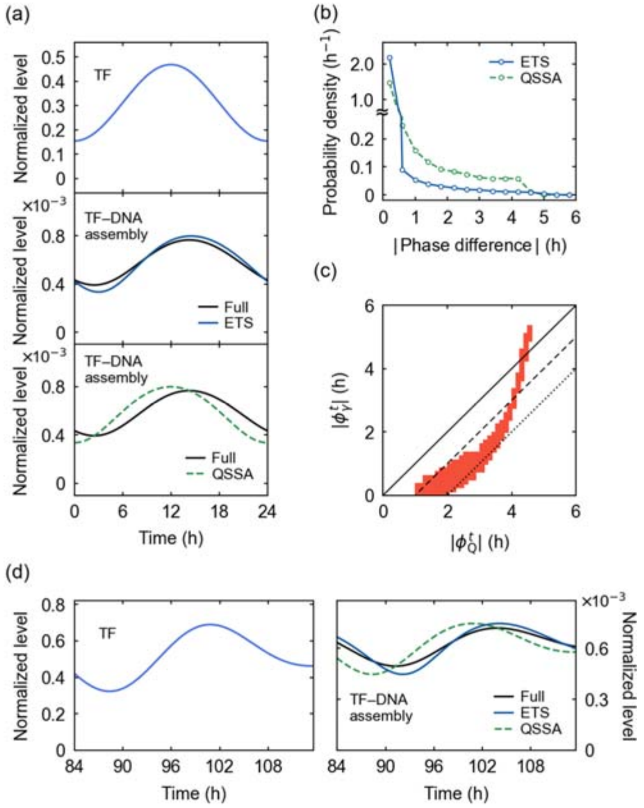


**Fig. S5. TF–DNA interaction modeling.** (a) Example time series of TF level $\overline{A}_{\mathrm{TF}}(\tau)$ in Eq. (S43) at the top, the full model-based TF–DNA assembly level $\overline{C}_{\mathrm{TF}}(\tau)$ and the ETS at the center, and $\overline{C}_{\mathrm{TF}}(\tau)$ and the QSSA at the bottom. $t = k_{\delta}^{-1}\tau$ as defined under Eq. (S20). (b) Probability distributions of $|\phi_{\gamma}^{t}|$ ("ETS") and $|\phi_{\mathrm{Q}}^{t}|$ ("QSSA") over randomly-sampled parameter sets in Table S3. (c) Scatter plot of $|\phi_{\mathrm{Q}}^{t}|$ and $|\phi_{\gamma}^{t}|$ when $|\phi_{\mathrm{Q}}^{t}|$ or $|\phi_{\gamma}^{t}|$ ≥1 h with randomly-sampled parameter sets in Table S3. A solid diagonal line corresponds to $|\phi_{\gamma}^{t}| = |\phi_{\mathrm{Q}}^{t}|$, a dashed diagonal line to $|\phi_{\gamma}^{t}| = |\phi_{\mathrm{Q}}^{t}| - 1\mathrm{h}$, and a dotted diagonal line to $|\phi_{\gamma}^{t}| = |\phi_{\mathrm{Q}}^{t}| - 2\mathrm{h}$. (d) Example time series of irregularly oscillating TF level $\overline{A}_{\mathrm{TF}}(\tau)$ in Eq. (S44) on the left and the full model-based TF–DNA assembly level $\overline{C}_{\mathrm{TF}}(\tau)$, the ETS, and the QSSA on the right. For more details of (a)–(d), refer to Text S8 and Tables S14 and S15.

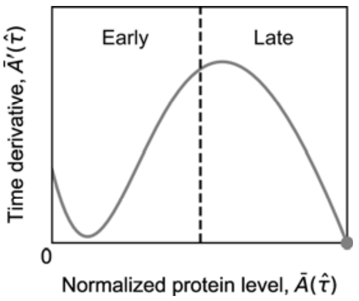


**Fig. S6. Phase portrait of induction kinetics with $\eta > \eta_c$ in the case of positive autoregulation.** A vertical dashed line splits the early and late stages of protein growth. A stable fixed point is indicated by a filled circle. For more details, refer to Text S9 and Table S8.

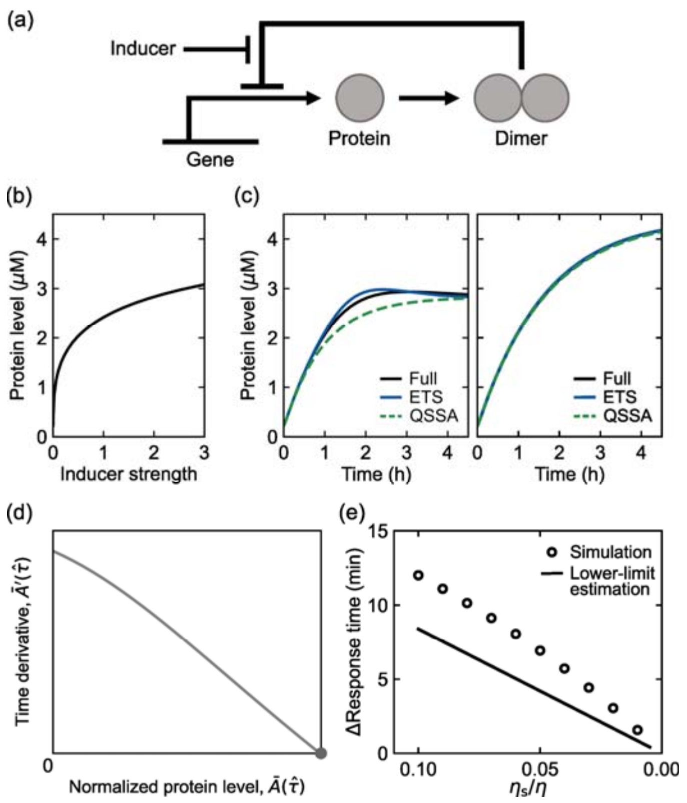


**Fig. S7. Negative autoregulation and induction kinetics.** (a) Protein production mechanism with negative autoregulation in the presence of inducers. (b) Bifurcation diagram of the simulated protein level as a function of $\eta$ (proxy for an inducer level). (c) Time-series of protein levels from the full, ETS, and QSSA models upon acute induction at time $0$ h with $\eta = 2$ (left) or $\eta = 80$ (right). The $\eta$ values were chosen for similar steady states to Fig. 2(c). (d) Phase portrait of induction kinetics. The stable fixed point is indicated by a filled circle. (e) The QSSA-to-full model difference in response time as a function of $\eta_s/\eta$ for $\eta \gg \eta_s$. Both the simulated difference and its analytically-estimated lower limit are presented. For more details of (b)–(e), refer to Text S10 and Table S16.

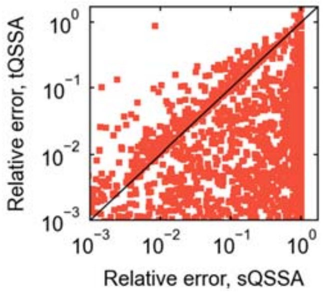


**Fig. S8. The sQSSA- and tQSSA-based parameter estimation.** The scatter plot of the relative errors of the sQSSA- and tQSSA-estimated $K$ values for a protein–protein interaction model. A diagonal line corresponds to the cases where the two estimates have the same relative errors. Most of the sQSSA-based estimates (77.4%) exhibit larger relative errors than the tQSSA-based ones, demonstrating the sQSSA's poorer parameter estimation. A subset of simulated conditions gave relative errors outside the presented ranges here, but they did not alter the observed tendency. For more details, refer to Text S12.

**Table S1. Enzyme–substrate pairs of metabolic reactions in *E. coli* (refer to Text S6).** $\varepsilon_1$, $\varepsilon_2$, $\varepsilon_{\mathrm{tQ}}$, and $\varepsilon_\gamma$ stand for $\max_\tau[\varepsilon_1(\tau)]$, $\max_\tau[\varepsilon_2(\tau)]$, $\max_\tau[\varepsilon_{\mathrm{tQ}}(\tau)]$, and $\max_\tau[\varepsilon_\gamma(\tau)]$ with known enzyme and substrate concentrations ("enzyme conc." and "substrate conc."), calculated by Eqs. (S29), (S31), (S32), and (S34). In this calculation, we applied rough approximations $C(t) \sim C_{\mathrm{tQ}}(t)$, $C_{\mathrm{tQ}}\left\{t - \left[k_\delta \Delta_{\mathrm{tQ}}(t)\right]^{-1}\right\} \sim C_{\mathrm{tQ}}(t)$, and $k_\delta \sim r_{\mathrm{c}}$. The enzyme and substrate concentrations and kinetic parameters were collected from Refs. [S26–S28], respectively, except the oxaloacetate concentration [S29].

| Enzyme | Substrate | Enzyme conc. (mM) | Substrate conc. (mM) | $K$ (mM) | $r_{\mathrm{c}}$ (ms$^{-1}$) | $\varepsilon_1$ | $\varepsilon_2$ | $\varepsilon_{\mathrm{tQ}}$ | $\varepsilon_\gamma$ |
|---|---|---|---|---|---|---|---|---|---|
| 6-Phosphofructokinase | Fructose 6-phosphate | 0.01 | 8.8 | 0.16 | 0.17 | $3.6\times10^{-10}$ | $1.9\times10^{-5}$ | $3.5\times10^{-7}$ | $6.5\times10^{-12}$ |
| Arginine decarboxylase | Arginine | 0.002 | 0.6 | 0.65 | 0.08 | $3.2\times10^{-7}$ | $3.9\times10^{-4}$ | $4.4\times10^{-4}$ | $1.7\times10^{-7}$ |
| Aspartate ammonia-lyase | Aspartate | 0.08 | 4.2 | 1.2 | 0.18 | $8.3\times10^{-6}$ | 0.003 | $7.3\times10^{-4}$ | $1.8\times10^{-6}$ |
| Aspartate carbamoyltransferase | Aspartate | 0.08 | 4.2 | 12.4 | 0.42 | $3.6\times10^{-6}$ | $9.5\times10^{-4}$ | 0.003 | $2.7\times10^{-6}$ |
| Aspartate transaminase | Aspartate | 0.02 | 4.2 | 4.0 | 0.53 | $8.6\times10^{-7}$ | $6.6\times10^{-4}$ | $6.3\times10^{-4}$ | $4.2\times10^{-7}$ |
| Glutamate decarboxylase | Glutamate | 0.15 | 96.0 | 2.32 | 0.02 | $1.3\times10^{-9}$ | $3.6\times10^{-5}$ | $8.8\times10^{-7}$ | $3.2\times10^{-11}$ |
| Ornithine decarboxylase | Ornithine | $5.9\times10^{-4}$ | 0.01 | 3.3 | 0.003 | $9.6\times10^{-11}$ | $5.4\times10^{-7}$ | $1.8\times10^{-4}$ | $9.6\times10^{-11}$ |
| Phosphoenolpyruvate carboxylase | Phosphoenolpyruvate | 0.007 | 0.18 | 0.19 | 0.54 | $5.0\times10^{-5}$ | 0.005 | 0.005 | $2.5\times10^{-5}$ |
| Succinate dehydrogenase | Succinate | 0.09 | 0.57 | 0.002 | 0.09 | $5.9\times10^{-7}$ | $7.6\times10^{-4}$ | $3.2\times10^{-6}$ | $2.4\times10^{-9}$ |
| Succinate dehydrogenase | Fumarate | 0.09 | 0.12 | 0.005 | 0.002 | 0.03 | 0.08 | 0.02 | 0.002 |
| Malate dehydrogenase | Oxaloacetate | 0.09 | $4.9\times10^{-4}$ | 0.04 | 0.93 | $4.1\times10^{-4}$ | $2.3\times10^{-4}$ | 0.40 | 0.11 |

**Table S2. The PTS system of *E. coli* (refer to Text S6).** The cellular concentration of the PTS subunit IICB (transporter) is 0.025 nmol/mg cell dry weight [S30] and we considered 0.76 mg/L initial cell dry density in a culture medium [S31] to calculate the transporter concentration here. Glucose (nutrient) concentration was obtained from Ref. [S31] and $K$, $r_c$, and $k_\delta$ from Ref. [S30]. We used a cell growth rate of 0.89 $h^{-1}$ [S31]. $\varepsilon_1$, $\varepsilon_2$, $\varepsilon_{tQ}$, and $\varepsilon_\gamma$ stand for $\max_\tau[\varepsilon_1(\tau)]$, $\max_\tau[\varepsilon_2(\tau)]$, $\max_\tau[\varepsilon_{tQ}(\tau)]$, and $\max_\tau[\varepsilon_\gamma(\tau)]$ with known transporter and nutrient concentrations, calculated by Eqs. (S29), (S31), (S32), and (S34). In this calculation, we applied rough approximations $C(t) \sim C_{tQ}(t)$ and $C_{tQ}\left\{t - \left[k_\delta \Delta_{tQ}(t)\right]^{-1}\right\} \sim C_{tQ}(t)$.

| Transporter conc. (mM) | Nutrient conc. (mM) | $K$ (mM) | $r_c$ ($ms^{-1}$) | $k_\delta$ ($ms^{-1}$) | $\varepsilon_1$ | $\varepsilon_2$ | $\varepsilon_{tQ}$ | $\varepsilon_\gamma$ |
|---|---|---|---|---|---|---|---|---|
| $1.9\times10^{-8}$ | 11.1 | 0.02 | 0.08 | 0.09 | $8.8\times10^{-18}$ | $2.8\times10^{-12}$ | $5.1\times10^{-9}$ | $1.3\times10^{-17}$ |

**Table S3. Parameter ranges of protein–protein and TF–DNA interaction models (refer to Texts S7 and S8).** Columns "PPI" and "TDI" indicate the relevance to the protein–protein and TF–DNA interaction models, respectively. $T = 24$ h. *The parameter is sampled uniformly on a logarithmic scale to cover a wide range of the order of magnitude. **It is sampled to assign $K$ in Eq. (S21) by dividing the sampled value by $\overline{A}_{\max}$. †The value is relevant to the TF–DNA interaction model: the minimum value is set to ensure the practically-meaningful oscillation of $\bar{C}_{\mathrm{TF}}(\tau)$ and the maximum value is set to avoid any instance of $\bar{A}_{\mathrm{TF}}(\tau) = 0$ for the condition of little stochasticity in $A_{\mathrm{TF}}(t)$ (Text S4). For the practically-meaningful oscillation of $\bar{C}(\tau)$ in the case of protein–protein interactions, we consider only the parameters of $\alpha_{\mathrm{C}} \geq 0.2$ where $\alpha_{\mathrm{C}}$ denotes the peak-to-trough difference of $\bar{C}(\tau)$ divided by the peak level. ††After the sampling of the value, we only consider the case $A_{\mathrm{TF}}(t)V \geq 10$ to satisfy the condition of little stochasticity in $A_{\mathrm{TF}}(t)$ (Text S4).

| Parameter | PPI | TDI | Minimum | Maximum | Remarks |
|---|---|---|---|---|---|
| $\overline{A}_{\max}$* | ✓ | ✓ | 0.01 | 100 | Inferred from Refs. [S4,S32]. |
| $\overline{B}_{\max}$* | ✓ | | 0.01 | 100 | The same range as $\overline{A}_{\max}$. |
| $K\overline{A}_{\max}$ (nM)** | | ✓ | 1 | 10 | Based on the rough range of clock protein levels in Refs. [S32,S33]. |
| $\alpha_{\mathrm{A}}$ | ✓ | ✓ | 0 (0.2†) | 1 (0.9†) | |
| $\alpha_{\mathrm{B}}$ | ✓ | | 0 | 1 | |
| $k_{\delta}$ ($h^{-1}$)* | ✓ | ✓ | 0.1 | 10 | Inferred from Ref. [S4]. |
| $\varphi_{\mathrm{B}}$ | ✓ | | 0 | $\pi$ | |
| $V$ ($nM^{-1}$)†† | | ✓ | 10 | 100 | Based on Ref. [S13]. |

**Table S4. Parameter ranges for ZTL profile simulation (refer to Text S7).** We narrowed down the initial parameter ranges from the sources listed here, to focus on the regime where at least the full model-based $A_s(t)$ reasonably approximates the experimental ZTL profile. *Relevant to a light condition. †Relevant to a dark condition. **The sampled parameter value for a light condition was used as the lower bound of the dark condition parameter.

| Parameter | Minimum | Maximum | Remarks |
|---|---|---|---|
| $g_A$ (nM·h$^{-1}$) | 0.5 | 2.5 | Inferred from simulated $A(t)$ belonging to the range of protein levels in Table S3 [S34]. |
| $r_A$ (h$^{-1}$) | 4.0 | 10.0 | Chosen to satisfy $r_c < r_A$ (from ZTL stabilization by GI) [S16–S18] and roughly based on the order of magnitude of very high protein degradation rates. |
| $r_c$ (h$^{-1}$) | 0.1 | 0.2 | Inferred from Ref. [S35] and chosen to satisfy $r_c < r_A$ from ZTL stabilization by GI [S16–S18]. |
| $k_d$ (h$^{-1}$)* | 0.1 | 0.5 | Derived from the range of $k_\delta$ in Table S3. |
| $k_d$ (h$^{-1}$)† | $k_d$** | 0.5 | Chosen to satisfy $k_d{}^* \leq k_d{}^\dagger$ because blue light enhances the ZTL–GI interaction [S16]. |
| $K$ (nM)* | 0.02 | 0.5 | Derived from the range of $K$ determined in Table S3. |
| $K$ (nM)† | $K$** | 0.5 | Chosen to satisfy $K^* \leq K^\dagger$ because blue light enhances the ZTL–GI interaction [S16]. |
| $w_{GI}$ (nM) | 4.0 | 10.0 | Derived from the range of protein levels in Table S3. |
| $w_{ZTL}$ (nM) | 2.0 | 5.0 | Derived from the range of protein levels in Table S3. |

**Table S5. Parameter ranges for induction kinetics simulation [refer to Texts S9 and S10 (associated with Section *Autogenous control* in the main text)].** We randomly selected the parameter values from the ranges here and chose $s = 0.05a_0$ for $s \ll a_0$. We then only considered the parameter values with $A_2(t), A(t) - 2A_2(t) \gg V^{-1}$ (at least at the steady state) for the validity of Eqs. (S47) and (S48) [or (S68)] (Text S12). $^*K \equiv (k_\mathrm{d} + k_\mathrm{dlt} + r_\mathrm{c})/k_\mathrm{a}$ and $\widehat{K}_\mathrm{TF} \equiv (k_\mathrm{TFd} + k_\mathrm{dlt})/\widehat{k}_\mathrm{TFa}$ whereby $k_\mathrm{a}$ and $\widehat{k}_\mathrm{TFa}$ are calculated. $^{**}$The value is relevant to the model of negative autoregulation. $^\dagger$In the case of Eq. (S55), we only considered $b_0 = 100$ h$^{-1}$, $r_\mathrm{c} = 0$ h$^{-1}$, and the other sampled parameters with $10 \le \kappa B_0$ (and $\eta_\mathrm{s} < 10^4$ in the negative autoregulation case) to satisfy Eq. (S55) (and to cover $\eta \gg \eta_\mathrm{s}$ in the negative autoregulation case; Text S12). $^{\dagger\dagger}$The parameter is sampled uniformly on a logarithmic scale to cover a wide range of the order of magnitude.

| Parameter | Minimum | Maximum | Remarks |
|---|---|---|---|
| $\eta^{\dagger\dagger}$ | $\eta_\mathrm{c}$ ($10^{-6**}$) | $10^6$ | |
| $a_0$ (h$^{-1}$) | 90 | 500 | Inferred from Ref. [S36]. |
| $a_1$ (h$^{-1}$) | 50 | 300 | Inferred from Refs. [S37,S38]. |
| $b_0$ (h$^{-1}$)$^\dagger$ | 10 | 100 | Inferred from Ref. [S39]. |
| $k_\mathrm{dlt}$ (h$^{-1}$) | 0.2079 | 2.079 | Corresponding to 20–200-min bacterial doubling time. |
| $k_\mathrm{d}$ (h$^{-1}$)$^{\dagger\dagger}$ | 0.1 | 10 | The range of $k_\delta$ in Table S3. |
| $k_\mathrm{TFd}$ (h$^{-1}$)$^{\dagger\dagger}$ | 0.1 | 10 | The range of $k_\delta$ in Table S3. |
| $r_\mathrm{c}$ (h$^{-1}$)$^\dagger$ | 0 | 5 | Inferred from Ref. [S40]. |
| $K$ ($\mu$M)$^{*\dagger\dagger}$ | 0.0001 | 1000 | Derived from Refs. [S38,S41,S42]. |
| $\widehat{K}_\mathrm{TF}$ ($\mu$M)$^{*\dagger\dagger}$ | 0.0001 | 1000 | Derived from Refs. [S38,S41,S42]. |
| $V$ (nM$^{-1}$) | 0.1 | 1 | Inferred from Ref. [S42]. |

**Table S6. Parameter ranges for protein degradation simulation [refer to Text S11 (associated with Section *Rhythmic degradation of circadian proteins* in the main text)].** We randomly selected the parameter values from the ranges here with $n = 1, 2, 3$ and only considered the simulation results with periodic $A(t)$ of 1 nM $\leq \max_t[A(t)] \leq$ 10 nM as in Table S3. We chose $T =$ 24 h and $\alpha_g = 1$. $^*a_i = k_i B$ $(0 \leq i \leq n-1)$ and $k_i$ and $B$ denote the protein's $(i+1)$-th modification rate coefficient and the ubiquitin ligase or kinase concentration, respectively.

| Parameter | Minimum | Maximum | Remarks |
|---|---|---|---|
| $g_{\max}$ (nM·h$^{-1}$) | 0.1 | 3.0 | Inferred from simulated $A(t)$ belonging to the range of protein levels in Table S3 and inclined to have rhythmic $r(t)$ [S34]. |
| $k_i$ (nM$^{-1}$h$^{-1}$)$^*$ | 0.006 | 0.06 | Based on Ref. [S22]. |
| $B$ (nM)$^*$ | 10 | 100 | Roughly based on ubiquitin ligase and kinase levels in Refs. [S43,S44]. |
| $r_c$ (h$^{-1}$) | 0.5 | 5.0 | Based on Refs. [S22,S35]. |

**Table S7. Parameters used in Figs. 1(a) and 1(b).**

| Parameter | (a) | (b) |
|---|---|---|
| $k_\delta$ (h$^{-1}$) | 0.13 | 0.28 |
| $K$ (nM) | | 8.25 |
| $V$ (nM$^{-1}$) | | 48.41 |
| $\bar{A}_{\max}$ | 1.79 | 0.47 |
| $\bar{B}_{\max}$ | 0.52 | |
| $\alpha_A$ | 0.76 | 0.67 |
| $\alpha_B$ | 0.59 | |
| $\varphi_B$ | 0.59 | |
| $T$ (h) | 24 | 24 |

**Table S8. Parameters used in Figs. 2(b)–2(d) and S6.**

| Parameter | 2(b) | 2(c), left | 2(c), right | 2(d) | S6 |
|---|---|---|---|---|---|
| $\eta$ | | 2.42 | 200 | | 1.64 |
| $s$ (h$^{-1}$) | 23.37 | | | | |
| $a_0$ (h$^{-1}$) | 467.38 | | | | |
| $a_1$ (h$^{-1}$) | 297.98 | | | | |
| $b_0$ (h$^{-1}$) | 100 | | | | |
| $k_{\mathrm{dlt}}$ (h$^{-1}$) | 0.56 | | | | |
| $k_{\mathrm{d}}$ (h$^{-1}$) | 0.58 | | | | |
| $k_{\mathrm{TFd}}$ (h$^{-1}$) | 3.83 | | | | |
| $r_{\mathrm{c}}$ (h$^{-1}$) | 0 | | | | |
| $k_{\mathrm{a}}$ ($\mu$M$^{-1}$h$^{-1}$) | 0.019 | | | | |
| $\hat{k}_{\mathrm{TFa}}$ (nM$^{-1}$h$^{-1}$) | 0.37 | | | | |
| $V$ (nM$^{-1}$) | 0.53 | | | | |

**Table S9. Parameters used in Fig. 3(c).** $^*k_0 = 0.04$ nM$^{-1}$h$^{-1}$ (protein modification rate coefficient) and $B = 33.87$ nM (ubiquitin ligase concentration).

| Parameter | Value |
|---|---|
| $g_{\max}$ (nM·h$^{-1}$) | 2.98 |
| $\alpha_{\mathrm{g}}$ | 1 |
| $T$ (h) | 24 |
| $a_0$ (h$^{-1}$) | $k_0 B^*$ |
| $r_{\mathrm{c}}$ (h$^{-1}$) | 1.30 |

**Table S10. Parameters used in Figs. S1(a) and S1(b).** $T = 24$ h.

| Parameter | (a) | (b) |
|---|---|---|
| $K\overline{A}_{\max}$ (nM) | 5.86 | 4.2 |
| $K\overline{B}_{\max}$ (nM) | 14.63 | |
| $\alpha_{\mathrm{A}}$ | 0.61 | 0.71 |
| $\alpha_{\mathrm{B}}$ | 0.51 | |
| $\varphi_{\mathrm{B}}$ | 1.77 | |
| $V$ (nM$^{-1}$)$^{\dagger\dagger}$ | | 45.3 |

**Table S11. Parameters used in Fig. S3(a).**

| Parameter | Value |
|---|---|
| $k_\delta$ (h$^{-1}$) | 0.17 |
| $\bar{A}_{\max}$ | 1.79 |
| $\bar{B}_{\max}$ | 0.52 |
| $\alpha_{\mathrm{A}}$ | 0.76 |
| $\alpha_{\mathrm{B}}$ | 0.59 |
| $\varphi_B$ | 0.59 |
| $T$ (h) | 24 |

**Table S12. Parameters used in Fig. S3(f).** $N = 10$, $k_\delta = 0.18$ h$^{-1}$, and $T_{B,i} = T_{A,i}$.

| Parameter | Values for $\boldsymbol{i = 1, 2, \cdots, N}$ $\boldsymbol{(N = 10)}$ | | | | | | | | | |
|---|---|---|---|---|---|---|---|---|---|---|
| $\bar{A}_{\max,i}$ | 0.65 | 1.31 | 1.18 | 6.00 | 0.53 | 0.14 | 2.71 | 5.70 | 1.42 | 9.32 |
| $\bar{B}_{\max,i}$ | 0.13 | 5.38 | 0.31 | 0.45 | 0.13 | 0.47 | 9.07 | 0.25 | 3.51 | 5.60 |
| $\alpha_{\mathrm{A},i}$ | 1.00 | 0.73 | 0.76 | 0.57 | 0.72 | 0.60 | 0.60 | 0.58 | 0.86 | 0.97 |
| $\alpha_{\mathrm{B},i}$ | 0.50 | 0.75 | 0.93 | 0.51 | 0.52 | 0.87 | 0.87 | 0.52 | 0.65 | 0.76 |
| $\varphi_{\mathrm{A},i}$ | -2.68 | -1.02 | -2.69 | 2.59 | -2.56 | -2.60 | -0.75 | 1.16 | -2.12 | 0.29 |
| $\varphi_{B,i}$ | -0.07 | -2.94 | 2.51 | -0.20 | -1.26 | -2.37 | -2.58 | 0.72 | -3.07 | -2.92 |
| $T_{A,i}$ (h) | 12.9 | 12.0 | 27.7 | 14.2 | 29.2 | 38.1 | 32.3 | 32.5 | 13.6 | 27.9 |

**Table S13. Parameters used in Figs. S4(a) and S4(b).**

| Parameter | Value |
|---|---|
| $g_{\mathrm{A}}$ (nM·h$^{-1}$) | 2.17 |
| $r_{\mathrm{A}}$ (h$^{-1}$) | 9.23 |
| $r_{\mathrm{c}}$ (h$^{-1}$) | 0.11 |
| $k_{\mathrm{d}}$ in light (h$^{-1}$) | 0.159 |
| $k_{\mathrm{d}}$ in darkness (h$^{-1}$) | 0.163 |
| $K$ in light (nM) | 0.22 |
| $K$ in darkness (nM) | 0.24 |
| $w_{\mathrm{GI}}$ (nM) | 6.87 |
| $w_{\mathrm{ZTL}}$ (nM) | 3.04 |

**Table S14. Parameters used in Fig. S5(a).**

| Parameter | Value |
|---|---|
| $k_\delta$ (h$^{-1}$) | 0.28 |
| $K$ (nM) | 8.25 |
| $V$ (nM$^{-1}$) | 48.41 |
| $\overline{A}_{\mathrm{max}}$ | 0.47 |
| $\alpha_{\mathrm{A}}$ | 0.67 |
| $T$ (h) | 24 |

**Table S15. Parameters used in Fig. S5(d).** $N = 10$, $k_\delta = 0.19$ h$^{-1}$, $K = 6.63$ nM, and $V = 82.43$ nM$^{-1}$.

| Parameter | Values for $\boldsymbol{i = 1, 2, \cdots, N}$ $\boldsymbol{(N = 10)}$ | | | | | | | | | |
|---|---|---|---|---|---|---|---|---|---|---|
| $\overline{A}_{\mathrm{max},i}$ | 0.47 | 1.01 | 1.06 | 0.17 | 0.92 | 1.49 | 0.57 | 1.49 | 0.21 | 1.09 |
| $\alpha_{\mathrm{A},i}$ | 0.79 | 0.86 | 0.74 | 0.73 | 0.67 | 0.88 | 0.62 | 0.70 | 0.66 | 0.80 |
| $\varphi_{\mathrm{A},i}$ | 2.99 | 0.60 | 1.73 | 2.45 | -0.49 | 1.25 | 0.84 | 0.32 | 1.19 | -2.19 |
| $T_{\mathrm{A},i}$ (h) | 31.0 | 29.8 | 38.5 | 36.4 | 29.7 | 26.6 | 17.6 | 21.6 | 23.9 | 25.8 |

**Table S16. Parameters used in Figs. S7(b)–S7(e).**

| Parameter | (b) | (c), left | (c), right | (d) | (e) |
|---|---|---|---|---|---|
| $\eta$ | | 2 | 80 | 1.64 | |
| $s$ (h$^{-1}$) | 23.37 | | | | |
| $a_0$ (h$^{-1}$) | 467.38 | | | | |
| $a_1$ (h$^{-1}$) | 297.98 | | | | |
| $b_0$ (h$^{-1}$) | 100 | | | | |
| $k_{\mathrm{dlt}}$ (h$^{-1}$) | 0.56 | | | | |
| $k_{\mathrm{d}}$ (h$^{-1}$) | 0.58 | | | | |
| $k_{\mathrm{TFd}}$ (h$^{-1}$) | 3.83 | | | | |
| $r_{\mathrm{c}}$ (h$^{-1}$) | 0 | | | | |
| $k_{\mathrm{a}}$ ($\mu$M$^{-1}$h$^{-1}$) | 0.019 | | | | |
| $\hat{k}_{\mathrm{TFa}}$ (nM$^{-1}$h$^{-1}$) | 0.37 | | | | |
| $V$ (nM$^{-1}$) | 0.53 | | | | |
